# Chemogenetic activation of hypothalamic oxytocin neurons reorients face exploration toward the eyes in macaques

**DOI:** 10.64898/2026.09.17.751377

**Authors:** Antoine Ameloot, Eloïse Disarbois, Heba Elseedy, Coline Duperron, Valery Grinevich, Angela Sirigu, Jean-René Duhamel

## Abstract

Oxytocin (OT) shapes social behavior across species. However, in humans and non-human primates, its effects have been studied mainly through exogenous administration, despite uncertainty about the extent to which peripherally administered OT reaches the brain and limited control over the neural pathways it engages. To overcome these limitations, we developed a chemogenetic approach targeting the endogenous OT system by selectively activating OT neurons in the paraventricular nucleus (PVN) and supraoptic nucleus (SON) of macaques using excitatory DREADDs. We combined this approach with digit-tracking, a non-invasive, calibration-free assay of face exploration in freely moving animals. In this task, subjects explored blurred face images through a finger-controlled viewing window, providing a proxy for gaze exploration and enabling behavioral assessment of OT neuron activation. DREADD-mediated activation redirected ongoing face exploration toward the eyes at the expense of non-face regions, increasing the consistency of eye-directed exploration and the time spent inspecting the eyes once reached. Immunohistochemical analysis confirmed selective DREADD expression in OT neurons of the PVN and SON. These findings provide the first causal evidence in primates that hypothalamic OT neurons regulate social visual orienting by increasing exploration of the eyes, thereby linking a hallmark effect of intranasal OT administration to defined central OT pathways and establishing chemogenetic access to this system in freely behaving macaques.

## INTRODUCTION

Among social animals, the benefits of group living depend on the flexible adjustment of behavior to ongoing group dynamics^1^. In primates, navigating complex and evolving group structures requires efficient use of information about others to guide social decisions^2^. Facial features, especially the eyes, convey rich information about identity, internal states and intentions, placing their processing at the core of primate social cognition^3,4^. These capacities rely on distributed but partly specialized neural networks shaped by social evolution^5,6^. The oxytocin (OT) system is well positioned to modulate these networks through widespread OT receptor expression^7,8,9^ and the broad long-range projections of hypothalamic OT neurons^9,10,11,12^. Consistent with this anatomical organization, OT has been implicated in the enhancement of socially relevant cue processing and adaptive social behavior^13,14,15^.

In rodents and other non-primate animal models, invasive cellular and circuit-level approaches have enabled mechanistic dissection of OT pathways and their causal contributions to behavior^16,17^. These studies show that OT neurons are organized into anatomically defined populations^9,18^ with shared, distinct, or even opposing behavioral functions. For example, parvocellular OT neurons in the paraventricular nucleus (PVN) coordinate magnocellular OT populations in the PVN and supraoptic nucleus (SON) to promote social interaction^19^, whereas distinct OT populations in the bed nucleus of the stria terminalis and retrochiasmatic supraoptic nucleus contribute to social avoidance^20,21^. This functional heterogeneity highlights the importance of targeting defined OT populations.

By contrast, human OT research relies mainly on minimally or non-invasive approaches, including peripheral assays, genetic studies, neuroimaging and exogenous OT administration, providing predominantly associative rather than mechanistic insight^22^. Intranasal OT (IN-OT), the standard delivery method, has uncertain central penetration and distribution, lacks pathway specificity and produces variable behavioral effects across individuals and contexts, limiting reproducibility and translational potential^23,24^. These limitations leave a major gap in understanding the function of the OT system in humans, only partly addressed by the restricted translational relevance of non-primate models^25,26,27^. Although monkeys provide a crucial bridge between rodent and human research^28,29^, OT studies in non-human primates have largely relied on the same approaches used in human research^30^, emphasizing the need for more targeted methods^31^.

Recent chemogenetic advances in macaques now enable selective and reversible control of neuronal activity in genetically defined cell populations^32^. Within the central OT system, neurons of the PVN and SON constitute the principal hypothalamic OT populations and give rise to the broadest projections and functional influence described so far^9,14,15,18^. These populations are functionally coupled^19,33,34,35^ and have previously been successfully targeted using DREADD-mediated activation in rodents^36,37^. Extending this approach to macaques therefore offers the opportunity to causally investigate how hypothalamic OT neurons regulate social behavior in a primate model relevant to human social cognition.

A key step in validating this approach is to determine whether chemogenetic activation of OT neurons can reproduce a hallmark behavioral effect repeatedly associated with exogenous OT administration. IN-OT has frequently been reported to increase attention to faces and, in particular, to the eyes in healthy adults, several clinical populations, and multiple primate species including macaques, marmosets, and bonobos^38,39,40,41,42,43,44,45,46^. However, reported effects vary widely across studies. Changes have been observed in different gaze metrics, restricted to specific stimulus categories or participant subgroups, or dependent on additional pharmacological manipulations such as opioid receptor blockade^47^. Other studies report unstable, null or even opposite effects on face exploration^48,49,50,51,52,53^. These inconsistencies likely arise, at least in part, from the pharmacokinetic and targeting limitations of intranasal delivery, leaving the mechanisms by which OT influences face exploration poorly understood. In particular, it remains unknown whether hypothalamic OT neuron activity causally regulates face exploration in primates, and whether this circuitry underlies the changes in social gaze variably attributed to IN-OT administration.

This study investigated how chemogenetic activation of OT neurons in the PVN and SON influences face exploration strategies in macaques viewing conspecific and human faces. Exploration was measured in home cages using digit-tracking, a calibration-free and minimally constraining method, already shown as a proxy for gaze exploration validated in freely behaving animals^54^. Activation of OT neurons redirected exploration toward the eyes at the expense of non-face regions, whereas the effects at the nose and mouth regions varied across subjects. This eyes-reorienting behavior was not found before viral delivery during pre-operative control sessions, indicating an OT-dependent effect rather than a DREADD-ligand artifact. Histological analysis further confirmed selective DREADD expression in OT neurons of the PVN and SON, providing definitive causal evidence that endogenous hypothalamic OT neurons regulate social visual orienting in primates by directing attention to socially relevant facial cues.

## RESULTS

### Experimental design and viral targeting

A face exploration task was used to test whether chemogenetic activation of OT neurons in the PVN and SON modulates ongoing spontaneous visual exploration behavior in macaques, with the prediction that it would increase exploration of the eye region, as previously reported after OT inhalation.

Monkeys performed the task from their home cages using a touchscreen device equipped with liquid reward delivery (Figure 1A). Visual exploration was measured with digit-tracking, a calibration-free and minimally constraining method validated in humans^55^ and in macaques^54^. In this paradigm, subjects explored blurred face images through a local clear aperture revealed by touch. Finger movements therefore provided a proxy for gaze exploration and a functional analogue of foveated visual sampling. The trajectory of the aperture was recorded during a 4 s free exploration period triggered by the first contact with the image (Figure 1B). Stimuli consisted of macaque and human faces varying in gaze direction, sex, and familiarity. Faces were presented at different screen positions (left, center, right) to minimize central viewing bias. Behavioral effects were assessed by comparing vehicle and ligand sessions conducted in pairs. Complete sessions comprised 105 face trials, and session pairs were repeated until at least 315 face explorations were collected per condition (Figure 1D).

**Figure 1.**
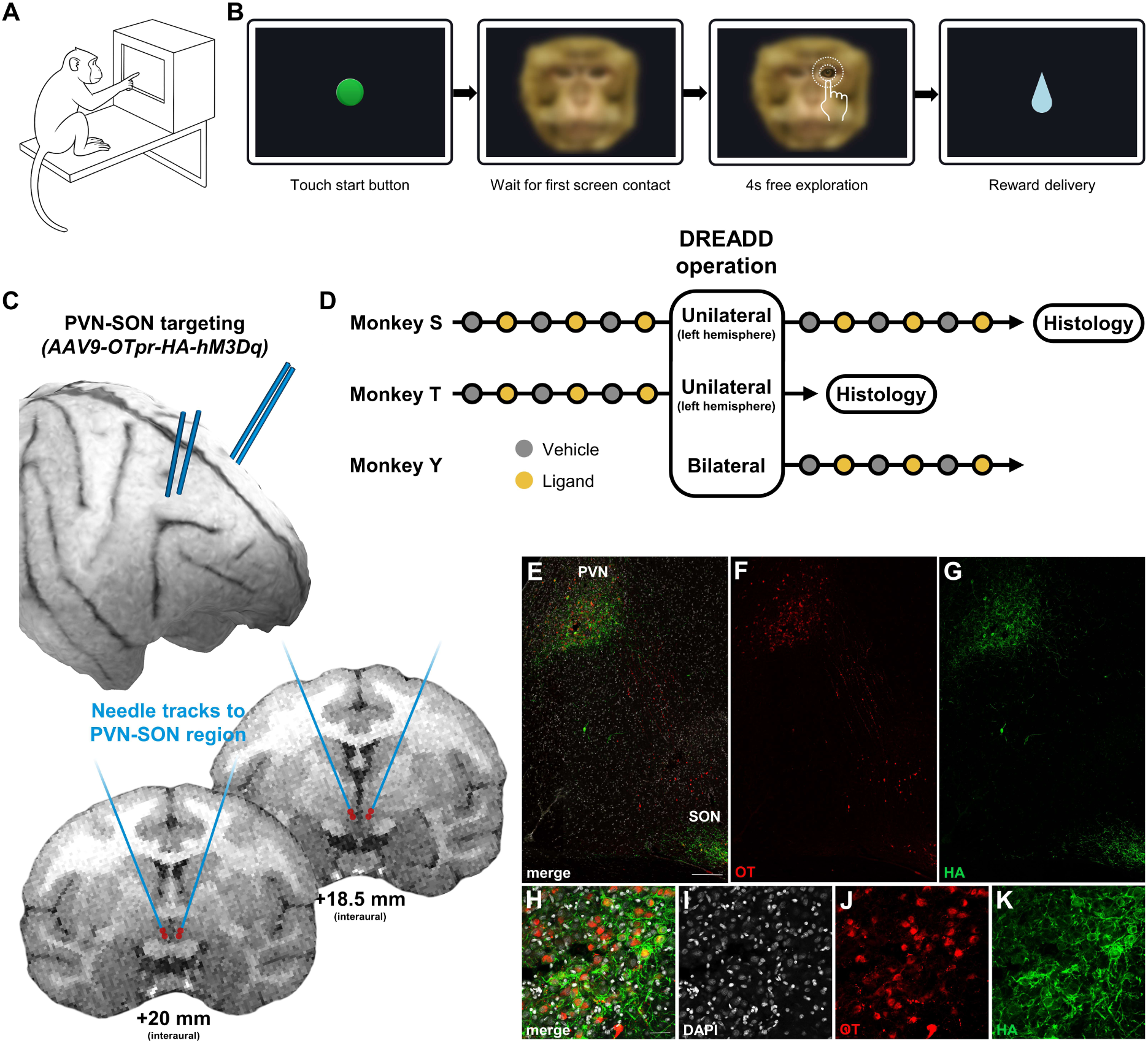
Face exploration task and DREADD viral targeting of PVN and SON OT neurons in macaques. **A.** Digit-tracking apparatus. Subjects interacted with a touch-sensitive screen placed against their home cages, displaying blurred face images. Screen contact revealed a clear area beneath the finger, providing a proxy of visual exploration. **B.** Task structure. Each trial began when the subject touched a start button to display a blurred face. The first screen contact triggered a 4 s free exploration window, followed by liquid reward delivery. **C.** Example 3D MRI reconstruction (top) and coronal sections (bottom) showing Brainsight-guided needle tracks and injection sites (red dots) in Monkey Y. Coronal sections are shown at AP +20.0 and +18.5 mm in subject space, corresponding approximately to Paxinos interaural levels +17.85 and +16.05 mm, respectively. **D.** Experimental timeline. Sessions (dots) are shown in chronological order across pre- and post-operative phases. Monkey S was tested with CNO (3 mg/kg), whereas Monkeys T and Y were tested with DCZ (0.1 mg/kg) as the DREADD ligand. Monkeys S and T received unilateral left-hemisphere viral vector injections, whereas Monkey Y received bilateral injections. **E–K.** Double immunofluorescence labeling of OT (red) and HA-tag (green) in Monkey T at an approximate Paxinos interaural level of +16.05 mm. **E–G.** Low-magnification images showing OT and HA-tag labeling in the PVN and SON, shown as a merged image in E and as single-channel images in F and G; scale bar, 200 µm. **H–K.** Higher-magnification images of the PVN showing OT, HA-tag, and DAPI labeling, shown as a merged image in H and as single-channel images in I–K; scale bar, 50 µm.

Monkey T was tested before unilateral AAV9-OTpr-HA-hM3Dq injection into the PVN-SON region, Monkey Y after bilateral injection, and Monkey S both before and after unilateral injection (Figures 1C and 1D; see Methods). Monkeys T and Y received the DREADD ligand DCZ, whereas Monkey S received CNO. Viral targeting was assessed by OT/HA-tag double immunofluorescence in the animals for which tissue was available, revealing robust and selective DREADD expression in OT neurons of the PVN and SON in Monkey T (Figures 1E–K) and Monkey S (Figures S1A–F).

### OT neuron activation biases face exploration toward the eyes

#### Spatial organization of exploration changes

We first examined OT-induced changes in face exploration in an ROI-free manner to characterize their spatial organization. Trial-by-trial exploration maps, representing the smoothed time spent at each image location to estimate foveal sampling, were compared between ligand and vehicle conditions using Cliff’s delta (δ), a non-parametric effect size reflecting the consistency of treatment-related differences across trials (see Methods). In both monkeys, high-δ clusters (top 2.5% of positive values) prominently overlapped the left eye, indicating a convergent increase in eye-directed exploration under OT neuron activation (Figures 2A and 2B, left panels). In Monkey S, positive values additionally extended across the eyes-nose region, suggesting a broader enhancement of exploration within the central facial area. Negative δ values were mainly located outside the principal facial features, although their spatial distribution differed between subjects. A decoding analysis applied to single-trial exploration maps significantly discriminated treatment condition in both monkeys (AUCs = 0.65, both *p* < 0.002, permutation test; Figures 2A and 2B, top right panels), indicating a modest but consistent exploration signature of OT neuron activation. Decoder weight maps highlighted the spatial features contributing most strongly to classification (Figures 2A and 2B, bottom right panels) and closely matched the corresponding δ maps (Monkey Y: *r* = 0.98; Monkey S: *r* = 0.96; both *p* < 0.001). Together, these complementary analyses indicate that OT neuron activation consistently biased exploration toward core facial features, particularly the eyes. Similar spatial signatures were observed when macaque and human faces were analyzed separately (Figure S2).

**Figure 2.**
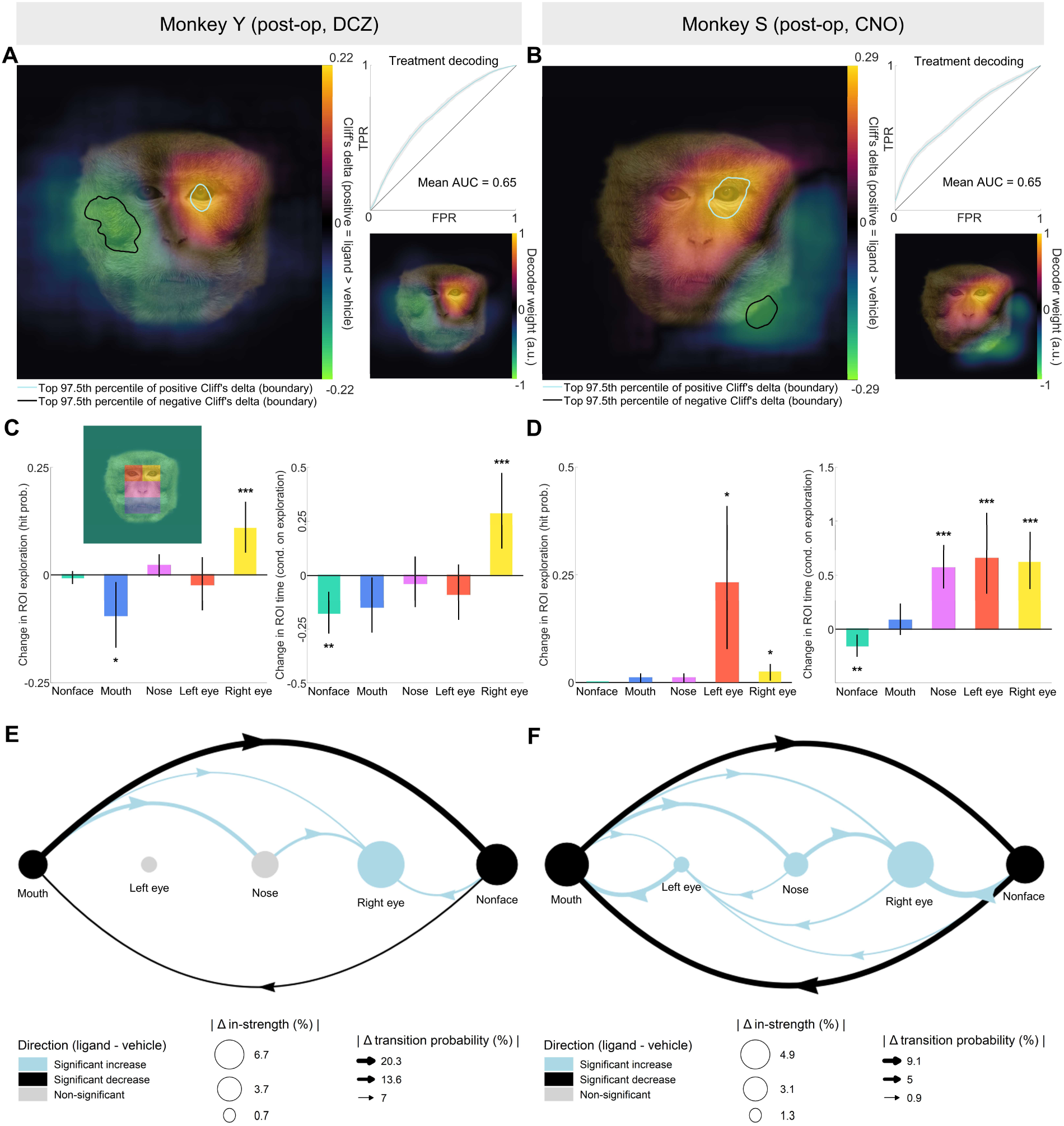
DREADD-mediated activation of PVN and SON OT neurons modulates face exploration. **A, B.** Spatially resolved analysis of exploration maps. Panels A and B correspond to Monkey Y and Monkey S. Left: Cliff’s delta (δ) map showing non-parametric effect sizes computed at each pixel across all ligand–vehicle trial pairs, reflecting the direction and consistency of treatment-related differences. Top right: ROC curve for treatment decoding using a logistic classifier (confidence intervals shown as gray shading). Bottom right: Decoder weight map highlighting informative spatial features. **C, D.** ROI exploration metrics estimated with a hurdle model (zero-inflated beta regression). Panels C and D correspond to Monkey Y and Monkey S. Left: relative change (ligand / vehicle − 1) in the probability that each ROI was explored at least once during the 4 s free exploration window (hit probability estimated from the zero-inflation part). Right: relative change (ligand / vehicle − 1) in the proportion of this 4 s window spent within an ROI when explored (estimated from the conditional beta part). Vertical lines indicate 95% confidence intervals; asterisks denote significance thresholds after Benjamini–Hochberg correction (*p < 0.05, **p < 0.01, ***p < 0.001). ROIs are illustrated on an example face image with colored areas (shown in panel C). **E, F.** ROI in-strength and transitions. Panels E and F correspond to Monkey Y and Monkey S. Graphs show differences (ligand − vehicle) in ROI in-strength (nodes, proportion of incoming transitions landing in each ROI) and transitions (edges, probability of transitioning from one ROI to another). In-strength significance was assessed by permutation testing, and transitions were estimated with a multinomial logistic model. For clarity, only significant edges after Benjamini–Hochberg correction are shown. Both monkeys were injected with AAV9 carrying the OTpr-HA-hM3Dq construct in the PVN-SON region. Monkey Y was tested with i.m. DCZ (0.1 mg/kg) and Monkey S with i.m. CNO (3 mg/kg) as DREADD ligands.

#### OT selectively enhances eye exploration

To quantify OT-DREADD effects on the exploration of specific facial features, behavior was analyzed within predefined ROIs corresponding to the eyes, nose, mouth, and non-face regions (Figure 2C, top). Because the maps revealed lateralized eye exploration, the eye ROI was further subdivided post hoc into left and right eyes. Exploration within each ROI was quantified using a hurdle-type model (see Methods), providing complementary estimates of hit probability (whether an ROI was explored at least once during the 4 s exploration window) and conditional proportion of time (proportion of exploration time spent within an ROI when explored). No significant interactions were observed between treatment and stimulus attributes (species, gaze direction, sex, or familiarity).

OT neuron activation consistently increased the probability of exploring the eyes (Figures 2C and 2D, left panels). In Monkey Y, the likelihood of exploring the left eye increased from 0.81 in vehicle to 0.89 in ligand trials (+10.9%, p < 0.001), whereas the right eye remained unchanged. In Monkey S, the largest increase was observed at the right eye (0.52 to 0.64, +23.1%, p = 0.012), with a smaller but significant increase at the left eye (0.97 vs 1.00, +2.3%, p = 0.048). This apparent discrepancy between ROI and δ-map analyses reflects ceiling effects and the different properties of the two measures. Left-eye exploration in Monkey S was already near ceiling under vehicle conditions (>97% of trials), limiting the relative increase in hit probability despite highly consistent exploration across trials. In contrast, the lower baseline probability at the right eye allowed a larger relative increase but produced weaker δ values because many trials still lacked exploration in both conditions. Thus, the two analyses capture complementary aspects of the same underlying effect and converge in identifying the eyes as the primary target of OT modulation. Effects at other facial regions were weaker and less consistent. Mouth exploration decreased in Monkey Y but remained unchanged in Monkey S, whereas nose and non-face regions showed no reliable changes in exploration probability.

OT neuron activation also increased the time spent exploring the eyes once visited, while reducing time allocated to non-face regions (Figures 2C and 2D, right panels). In Monkey Y, the proportion of the 4 s exploration window spent at the left eye increased from 0.037 to 0.047 (+28.9%, *p* = 0.001), whereas no reliable change was observed at the right eye. In Monkey S, exploration time increased at both eyes (right: +66.2%, *p* < 0.001; left: +61.6%, *p* < 0.001). Beyond the eyes, exploration time at the nose increased selectively in Monkey S (+56.6%, *p* < 0.001), whereas time spent in non-face regions decreased in both monkeys (Monkey Y: −17.8%, *p* = 0.003; Monkey S: −16.0%, *p* = 0.007). No reliable changes were observed at the mouth.

Overall, these results indicate that OT neuron activation increased both the likelihood and duration of eye exploration while reducing engagement with non-face regions.

#### Exploration flow dynamics

Next, how OT neuron activation reshaped transitions between facial regions beyond time-averaged spatial measures was examined using two complementary transition-based analyses. In-strength quantified the proportion of transitions landing in each ROI, providing a measure of how strongly each region attracted exploration. First-order transition probabilities characterized the pathways linking ROIs.

Across subjects, OT neuron activation increased in-strength at the eyes while reducing it at mouth and non-face regions (Figures 2E and 2F). In Monkey Y, the proportion of transitions landing in the left eye increased by 6.7 percentage points (pp), from 11.9% in vehicle to 18.6% in ligand trials (*p* = 0.005), whereas no reliable change was observed at the right eye. In Monkey S, in-strength increased at both eyes (right: +1.3 pp, *p* < 0.001; left: +4.9 pp, *p* < 0.001). Conversely, mouth and non-face regions showed reduced in-strength in both monkeys, indicating a redistribution of exploration toward the eyes. A more limited increase at the nose was additionally observed in Monkey S.

Transition analyses further showed that increased eye-directed exploration was supported by enhanced inflows from nose, mouth, and non-face regions (Figures 2E and 2F). In both monkeys, transitions from non-face regions toward the eyes increased markedly, whereas transitions between mouth and non-face regions decreased. In Monkey S, increased inflows from the mouth also contributed to enhanced exploration of the nose.

Together, these analyses indicate that OT neuron activation reorganized exploration dynamics toward the eyes by increasing both their attractiveness as exploration targets and the probability of transitions from other facial and non-face regions. More broadly, ROI-free and ROI-based analyses converged to show that OT neuron activation biased exploration toward the eyes while reducing engagement with non-face regions.

### No ligand-induced enhancement of eye exploration before viral vector delivery

To determine whether the post-operative behavioral changes reflected chemogenetic activation of OT neurons rather than nonspecific pharmacological effects, the same analyses were performed before viral vector delivery.

In Monkey T, tested with DCZ, pre-operative analyses revealed a spatial exploration pattern distinct from the post-operative OT effect (Figures S3A, S3C, and S3E). δ maps and decoder weights emphasized the lower face and mouth region rather than the eyes, and ROI analyses showed reduced eye exploration together with increased mouth exploration. Exploration dynamics further confirmed this mouth-centered reorganization, with increased inflows toward the mouth and reduced transitions toward the eyes.

In Monkey S, tested with CNO, pre-operative effects were weak and inconsistent (Figures S3B, S3D, and S3F). δ values were small in magnitude, decoding performance was near chance level (AUC = 0.54), and ROI analyses revealed no reliable changes in eye exploration. Minor transition effects were observed toward the mouth region but remained limited in magnitude.

Together, these pre-operative analyses show that DCZ and CNO per se did not reproduce the post-operative enhancement of eye exploration. When ligand effects were present, they were heterogeneous and largely opposite to the post-operative OT-related effects, indicating that the eye-focused reorganization observed after viral vector delivery reflects DREADD-mediated activation of OT neurons rather than nonspecific ligand effects.

### Sources of the rightward exploration bias

Exploration analyses revealed a rightward bias in the subject’s visual field (corresponding to the left side of the face stimulus), most evident as increased sampling of the left eye under OT (Figure 2). To clarify the origin of this asymmetry, and because a similar bias was observed prior to viral delivery (Figure S3), subsequent analyses examined whether it reflected motor asymmetry, ligand effects, or OT-dependent mechanisms.

Lateralized exploration can arise from different reference frames. A purely egocentric bias, for example due to motor asymmetries, should remain anchored to screen coordinates and therefore produce a stronger screen-referenced than face-referenced deviation. Conversely, a stimulus-centered bias, reflecting preferential exploration of one side of the face, should remain anchored to facial coordinates and produce a stronger face-referenced than screen-referenced deviation. Because faces were presented at multiple screen locations, these possibilities could be dissociated. To quantify exploration biases, we computed two lateralization indices (LIs): a screen-referenced LI and a face-referenced LI (Figure 3A).

**Figure 3.**
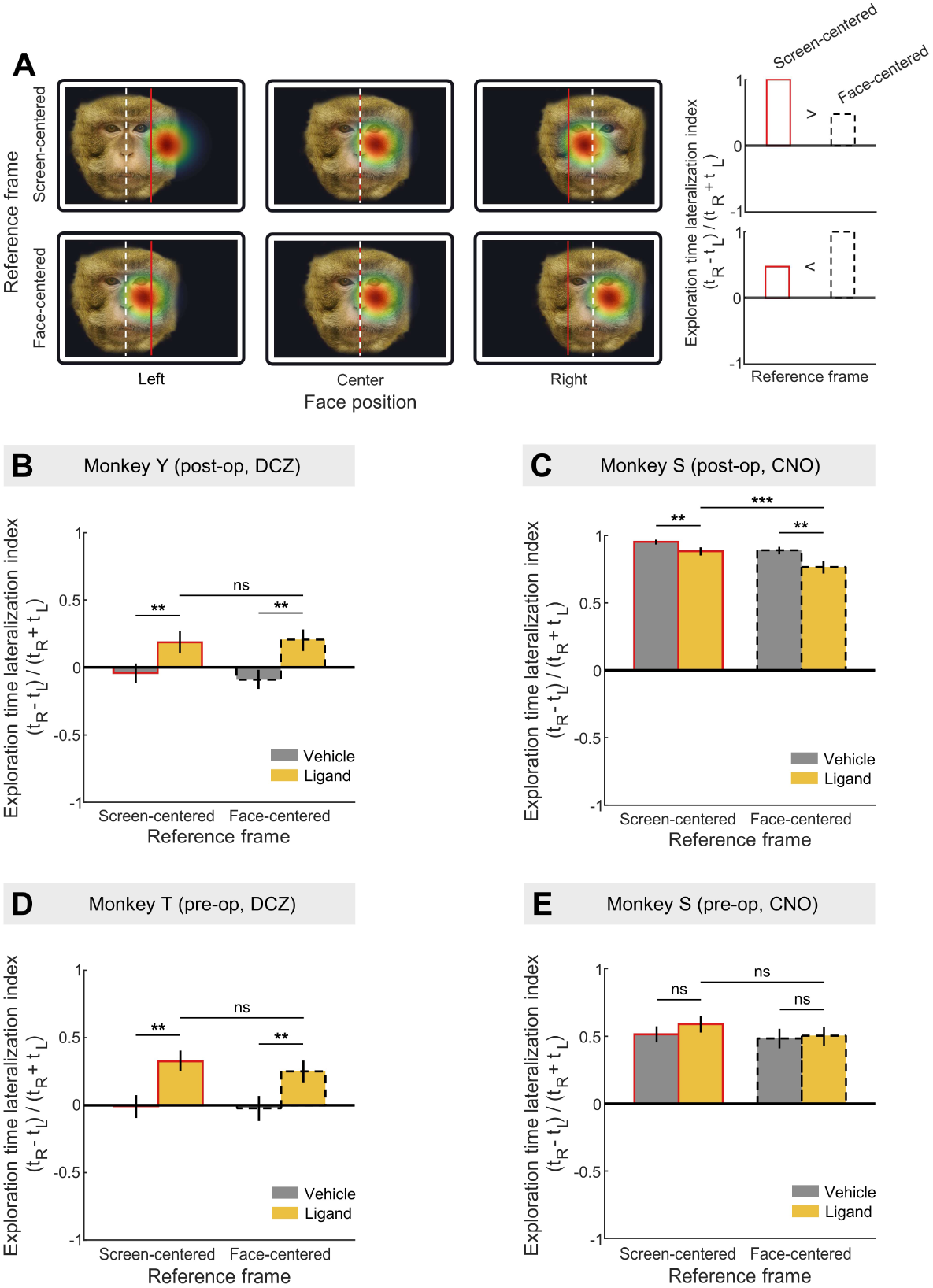
Lateralization index (LI) analysis across subjects. **A.** Conceptual distinction between screen-centered and face-centered lateralization. Simulated exploration maps are shown for faces presented at left, center, and right screen positions. Solid red and dashed white vertical lines indicate the screen and face midlines, respectively. Top row: idealized screen-centered bias, predicted to yield a larger screen-referenced than face-referenced lateralization index (LI). Bottom row: idealized face-centered bias, predicted to yield a larger face-referenced than screen-referenced LI. Bar graphs summarize the expected LI relationships under each hypothesis. **B–E**. Each panel corresponds to a subject in a specific experimental condition, as indicated by the panel titles. Within each panel, screen- and face-referenced LIs are shown for vehicle and ligand conditions. These indices quantify the normalized difference between time spent exploring the right and left sides of the reference midline. Values range from −1 (exclusive left-side exploration) to +1 (exclusive right-side exploration). Vertical lines indicate 95% confidence intervals; asterisks denote significance thresholds after Benjamini–Hochberg correction (*p < 0.05, **p < 0.01, ***p < 0.001).

Under vehicle conditions, exploration was symmetric in both Monkey Y and Monkey T, with no bias in either reference frame, ruling out a baseline motor asymmetry linked to handedness (Figures 3B and 3D). In contrast, DCZ induced a robust rightward shift in both monkeys, reflected in increases in both screen- and face-referenced LIs. Crucially, this effect was observed both before and after viral delivery, indicating that it does not depend on DREADD expression. Moreover, the bias was expressed across both reference frames, with no significant difference between screen- and face-referenced LIs (Monkey Y: *p* = 0.63; Monkey T: *p* = 0.11). This pattern argues against a purely egocentric motor bias, which would predict stronger screen-centered lateralization, as well as against a strictly stimulus-centered attentional effect, which would predict stronger face-referenced lateralization. Instead, the results indicate that the DCZ-induced rightward shift reflects a combination of egocentric and stimulus-centered influences. Together, these results indicate that DCZ induces a rightward shift in exploration that is independent of OT neuron activation and reflects a ligand-driven bias. In Monkey S, exploration under vehicle showed a strong rightward bias across reference frames, consistent with a pre-existing motor asymmetry (Figures 3C and 3E). Thus, while DCZ induces a rightward shift in face exploration in a DREADD-independent manner, CNO did not produce a comparable bias and instead reduced a pre-existing motor asymmetry following OT neuron activation, highlighting distinct ligand-specific effects.

### Cross-subject convergence of treatment effects

Given the small number of subjects and heterogeneous single-subject effects, additional analyses assessed whether treatment-related changes in exploration converged on shared spatial patterns across monkeys. To test this possibility, cross-subject decoding analyses were performed between Monkey Y (DCZ) and Monkey S (CNO) after vector delivery. Classifiers trained on one subject and tested on the other showed a trend toward above-chance performance (AUC = 0.61, *p* = 0.08; Figure S4A), indicating partial generalization of treatment-related exploration patterns across subjects. Spatial analyses revealed convergence centered on the left eye, whereas regions on the right side of the face showed more variable or divergent effects. ROI-based decoding confirmed this pattern (AUC = 0.63, *p* = 0.008), identifying the left eye as the most consistent feature supporting cross-subject generalization.

To determine whether this convergence could be explained by ligand effects alone, the same analysis was applied before vector delivery. Cross-subject decoding between Monkey T (DCZ) and Monkey S (CNO) showed no reliable generalization (AUC = 0.55, *p* = 0.16; Figure S4B), with weak and inconsistent spatial patterns, indicating that ligand effects per se do not account for the shared post-operative structure.

Next, ligand effects were compared before and after DREADD expression. In Monkey S, ROI-based decoding between pre- and post-operative sessions revealed a reversal of the treatment-related exploration pattern (AUC = 0.38, *p* = 0.015; Figure S4C), indicating that post-operative effects reflected a transformation of ligand-related influences rather than their persistence. Consistently, DCZ effects alone did not reproduce the post-operative pattern when comparing pre-operative Monkey T and post-operative Monkey Y (AUC = 0.49, *p* = 0.61; Figure S4D).

Together, these results indicate that chemogenetic activation of OT neurons induces a spatially consistent reorganization of face exploration across subjects, centered on the left eye, that cannot be explained by ligand effects alone.

## DISCUSSION

The present study investigated how chemogenetic activation of hypothalamic OT neurons influences face exploration in freely behaving macaques. In a home-cage digit-tracking task, activation of OT neurons in the PVN and SON consistently redirected exploration toward the eyes and away from non-face regions. Spatial mapping and ROI-based analyses converged in showing that OT neuron activation increased both the probability of exploring the eyes and the flow of exploration toward them from other facial and non-face regions. Although effects at the nose and mouth differed across subjects, these differences appeared to reflect subject-specific patterns superimposed on a common OT-driven shift toward the eyes. Pre-operative analyses further showed that DREADD ligands alone did not reproduce the post-operative enhancement of eye exploration, despite producing ligand-specific effects on exploration patterns. Histological analysis confirmed selective DREADD expression in OT neurons in the PVN and SON. Together, these findings provide causal evidence in primates that activation of hypothalamic OT neurons reshapes visual orienting toward socially informative facial cues.

The OT-driven increase in eye exploration was most strongly expressed at the left eye. However, several observations indicate that this lateralized component primarily reflected ligand-related and pre-existing motor influences rather than an intrinsic OT-specific bias. DCZ induced a rightward shift in exploration both before and after viral vector delivery, indicating a DREADD-independent effect. The DCZ-induced bias was expressed across both screen- and face-centered reference frames, indicating a combination of egocentric motor bias and stimulus-centered attentional effect.

The mechanisms underlying this DCZ-related effect remain uncertain. Although DCZ is a high-affinity DREADD agonist, it retains measurable off-target affinity at endogenous monoaminergic receptors at the dose used here^56^, and subtle behavioral effects have previously been reported in monkeys tested prior to viral vector delivery^57^. Such off-target influences could alter neural systems involved in spatial orienting or directional motor control, thereby biasing face exploration. By contrast, CNO did not induce a comparable rightward shift and instead reduced the strong baseline asymmetry observed in Monkey S following OT neuron activation. Despite these ligand-specific differences, both monkeys showed increased eye-directed exploration under OT activation, indicating that the core OT effect was not reducible to the lateralized bias itself. The interaction between OT effects and baseline lateralization may nevertheless have shaped how exploration was redistributed across the two eyes. In Monkey Y, OT-driven enhancement of eye exploration likely combined with the DCZ-related rightward bias, amplifying exploration of the left eye. In Monkey S, where baseline exploration was already strongly right-biased, OT activation appeared to broaden exploration toward both eyes in the absence of a CNO-driven directional effect. Because Monkey S received unilateral viral injection in the left hemisphere, however, a contribution of asymmetric OT activation to this redistribution cannot be excluded.

Although OT effects at the eyes and in non-face regions were broadly similar across monkeys, changes at the nose and mouth differed across subjects. These differences appear to reflect how a common OT-driven reorganization interacted with distinct baseline face-scanning strategies. Under vehicle conditions, Monkey S devoted a large proportion of exploration to non-face regions and the mouth, whereas Monkey Y distributed exploration more evenly across facial features, reflecting a more face-centered exploration pattern. Transition analyses indicated that OT neuron activation reshaped exploration flow in both monkeys by redirecting transitions from non-eye regions toward the eyes. In this reorganization, the nose tended to act as an intermediate relay, whereas the mouth became a less frequent endpoint of exploration. In Monkey S, where baseline exploration was dominated by non-face regions and the mouth, this redistribution increased transitions through the nose toward the eyes, plausibly contributing to the increased time spent at the nose under OT. In Monkey Y, where exploration was already more centered on facial features, the same reorganization primarily shifted exploration away from the mouth and toward the eyes.

Together, these findings suggest that activation of OT neurons does not impose a fixed exploration pattern but rather reshapes existing face-scanning strategies toward socially informative facial cues. This interpretation is consistent with evidence that socio-attentional phenotypes emerge from interactions between genetic, developmental, and experiential factors that shape face-processing circuits and their neuromodulatory systems^58,59,60,61,62,63^. OT signaling may therefore act less as a deterministic driver of social attention than as a modulatory system that amplifies endogenous biases toward socially relevant facial information.

The present analyses further suggest that OT neuron activation influenced multiple stages of face exploration. OT increased both the probability that exploration reached the eyes and the time spent exploring them once reached. Transition analyses showed that these effects were accompanied by a redistribution of exploration flows from non-eye regions toward the eyes. Increased transitions and hit probability are consistent with enhanced orienting toward socially relevant information, whereas prolonged exploration after arrival is more consistent with changes in the incentive or affective value of eye cues. Although the present data do not allow these processes to be dissociated, their coexistence argues against a purely attentional account and instead suggests that OT signaling influences both the selection and the evaluation of socially informative facial features.

The digit-tracking paradigm may be particularly sensitive to such effects because exploration requires active contact with facial features rather than passive observation. Direct gaze and eye cues can simultaneously provide valuable social information and elicit avoidance responses, creating an approach-avoidance conflict during face exploration. Consistent with this view, direct-gaze faces increase response latencies in macaques, suggesting that engagement with socially salient facial stimuli can carry an avoidance cost even in computerized tasks^64^. OT neuron activation may therefore promote eye exploration not only by increasing the value of eye cues but also by reducing reluctance to engage with them.

Several, non-exclusive mechanisms could contribute to these effects. OT signaling may increase the motivational value of social stimuli through actions on mesolimbic reward circuits, consistent with evidence that eye contact and mutual gaze engage reward-related networks in both humans and macaques^41,65,66,67^ and with rodent studies showing that PVN OT projections promote social motivation and reward^68,69,70^. OT signaling may also reduce the aversive salience of social stimuli through modulation of amygdala-centered circuits involved in social vigilance and threat processing. In both humans and macaques, IN-OT attenuates responses to threatening social cues and reduces vigilance toward socially aversive stimuli^39,71,72,73,74,75^, while studies in rodents demonstrate anxiolytic actions of PVN and SON OT pathways within amygdala networks^33,35,76^. Consistent with these effects, intra-amygdalar OT signaling has been shown to promote social investigation and other forms of social engagement in rodents^77,78,79,80^. Finally, OT may influence attentional orienting itself through actions on salience and arousal systems enriched in OT receptors in primates^7,26^. Rather than favoring one of these accounts, the present findings suggest that OT-dependent changes in face exploration emerge from coordinated modulation of distributed circuits involved in social valuation, emotional regulation, and attentional allocation.

A further limitation concerns the signaling mechanisms engaged by the present manipulation. Although chemogenetic activation of OT neurons evokes central and peripheral OT release in rodents^36,37^, these neurons can co-release glutamate and other neuromodulators^34,81^. The present findings therefore identify hypothalamic OT neurons as a causal substrate of eye-oriented face exploration, but do not establish which transmitter systems mediate this effect. Addressing this question will require combining cell-type-specific manipulations with physiological recordings and circuit-level interrogation of downstream targets in behaving primates.

Activation of hypothalamic OT neurons was sufficient to reproduce one of the behavioral effects most frequently associated with intranasal OT administration: increased exploration of the eyes. By linking this effect to selective activation of defined hypothalamic OT populations, the present findings provide causal support for the idea that endogenous OT circuitry contributes to the regulation of social visual orienting in primates. At the same time, the variability previously reported in IN-OT studies likely reflects the combined influence of pharmacokinetic limitations, species differences, individual variability, and differences across behavioral paradigms and social contexts^23,24^. Interestingly, increases in eye exploration after IN-OT are reported most consistently in paradigms using static face stimuli, whereas effects during naturalistic social interactions appear more variable in both humans and macaques^49,50,52^. Rather than reflecting a contradiction, these findings may indicate that OT consistently biases attention toward socially informative facial cues, while the behavioral consequences of this bias depend on the social context. The increased eye exploration observed during static face viewing may therefore reveal a core OT-dependent process that is expressed in more diverse and context-dependent ways during real social interactions.

Overall, this work provides the first causal evidence in primates that selective activation of hypothalamic OT neurons enhances attention to socially informative facial features. By enabling reversible and cell-type-specific manipulation of OT neurons in freely behaving macaques, this approach establishes a translational bridge between rodent circuit-level studies and human research that has relied largely on intranasal OT administration. More broadly, identifying central OT pathways that regulate eye engagement and social visual orienting may help inform future translational strategies targeting social-attentional dysfunctions associated with reduced eye contact and gaze avoidance, including autism spectrum disorder and social anxiety disorder^82,83,84^.

## MATERIALS AND METHODS

### Animals and ethics

Experimental procedures were authorized by the national ethics committee on animal experimentation N°42 and by the French Minister of Research and Innovation under the number APAFIS#30036-2021031218457217. All procedures conformed to European regulations and National Institutes of Health Guide for the Care and Use of Laboratory Animals, and ARRIVE guidelines. For this study, data were collected from three male long-tailed macaques at ISC-MJ: Monkey S (12 years, 9.7 kg), Monkey T (12 years, 9.5 kg), and Monkey Y (5 years, 4.5 kg). Animals were provided with a diet consisting of monkey chow, fresh fruits, and vegetables. Throughout the study, the minimum daily water intake for each animal was maintained at no less than 20 mL per kilogram of body weight, regardless of task performance, to ensure proper hydration and overall health. Each monkey had one day of free access to water each week. Enrichments were offered several times a week with different toys or substrates (boxes and puzzles containing dry fruit, swings) that promote play, curiosity, object manipulation, and foraging following recommendations from our own laboratory animal welfare committee.

### Viral vector production

An adeno-associated virus serotype 9 (AAV9) encoding HA-tagged hM3Dq under the control of a fragment of the murine oxytocin promoter was custom-produced by the Viral Vector Facility (University of Zurich and ETH Zurich). The recombinant vector (*ssAAV9/2-mOxy-HA_hM3D(Gq)-bGHp(A)*) was generated using a 2.6 kb promoter fragment previously validated for OT neuron targeting^33^ and the HA_hM3D(Gq) fragment from Addgene plasmid #50466. The final titer was 4.3 × 10¹³ vg/mL. Aliquots (50 µL) were stored at −80 °C until use.

### Surgical procedures and viral vector injections

Surgeries were performed under aseptic conditions. Anesthesia was induced with intramuscular ketamine (7 mg/kg) and medetomidine (50 µg/kg), then maintained with isoflurane (0.5-2%) via endotracheal intubation. Vital parameters were continuously monitored, and animals received antibiotic (amoxicillin, 20 mg/kg), anti-inflammatory (tolfenamic acid, 4 mg/kg), and analgesic (buprenorphine, 0.01 mg/kg) treatment during and after surgery.

Pre-operative MRI (3T) and CT scans were acquired at the CERMEP imaging facility (Bron, France) under anesthesia induced with intramuscular tiletamine-zolazepam (15 mg/kg). Images were co-registered and imported into Brainsight (Rogue Research, Canada) for trajectory planning and image-guided injections performed with the VetRobot arm, except for Monkey S, where coordinates were manually calculated from MRI and used for classical frame-based stereotaxic injection. A small craniotomy (1-2 mm) was made along the planned injection trajectory, and viral vector was infused through a 30-gauge needle guided by a 23-gauge outer needle. Two injections were made along each track, the first at the deepest site and the second 1 mm above. For the first site, the needle was lowered 200 µm past the target, immobilized for 2 min, then withdrawn to the injection position. The needle was kept stationary for 2 min before each infusion and for 5 min afterward; 4 µL was infused per site at 1 µL/min.

Monkeys S and T received unilateral viral vector injections, whereas Monkey Y received bilateral injections. In Monkey S, 4 injections were made along 2 tracks in the left hemisphere to target the PVN-SON region, with an anterior-mid site (AP 20.5 mm, DV 9.0 mm, ML −1.0 mm) and a mid-posterior site (AP 18.5 mm, DV 9.0 mm, ML −1.0 mm). Coordinates are expressed in millimeters relative to interaural zero in the macaque stereotaxic frame. Coordinates from Monkey S served as a reference for Brainsight-guided neuronavigation in the other monkeys, with final coordinates adjusted to each subject’s anatomy to target the same regions. Monkey T likewise received unilateral injections along 2 tracks in the left hemisphere targeting corresponding regions, but was euthanized 12 days after surgery following post-operative complications unrelated to the experimental procedures under investigation; only pre-surgical behavioral data and post-mortem histological data were therefore available for this animal. Monkey Y received bilateral injections, with 2 tracks per hemisphere targeting the same regions (trajectories shown in Figure 1C).

### Drug preparation and administration

For Monkey S, clozapine-N-oxide dihydrochloride (CNO; Bio-Techne, #6329) was administered intramuscularly at 3 mg/kg, using a solution freshly prepared by dilution in sterile saline. Deschloroclozapine (DCZ) was used in later experiments instead of CNO because of its higher potency and selectivity for DREADDs, allowing precise receptor activation at lower systemic concentrations^56^ and reducing the risk of CNO back-metabolism into clozapine and related off-target effects^85^. For the other monkeys, DCZ (Tocris, #7193) was administered intramuscularly at 0.1 mg/kg, a dose commonly used in NHP chemogenetic studies that produces high DREADD occupancy and is generally considered behaviorally inert prior to viral transduction^32,56,57^. DCZ was dissolved in DMSO, aliquoted and stored at −20 °C, then diluted in saline to a final DMSO concentration of 5% shortly before injection. The ligand or vehicle (saline for CNO; 5% DMSO in saline for DCZ) was injected 45 minutes before the behavioral task, which began at a fixed time to maintain consistency. The 45-min interval was selected based on previous studies demonstrating robust OT-neuron activation within this time window following DREADD ligand administration^36^. Ligand and vehicle sessions alternated, with at least one washout day between them.

### Histology

Histological analyses were performed in Monkeys S and T. Tissue from Monkey Y was not available because the animal remained in the colony after completion of the study. Animals were anesthetized with ketamine (10 mg/kg, i.m.) and euthanized with pentobarbital (1 mL/kg, i.v.), then perfused transcardially with ice-cold phosphate-buffered saline (PBS, 0.1 M, pH 7.4) followed by 4% paraformaldehyde (PFA) in PBS. Brains were extracted and post-fixed overnight in 4% PFA at 4 °C. Following rinsing in phosphate buffer (PB), tissues were cryoprotected in 20% sucrose in PB at 4 °C until they sank. Samples were then rapidly frozen on dry ice and sectioned coronally at 40 μm using a cryostat (Leica CM3050S). Sections were rinsed in PB, collected sequentially in an ethylene glycol-based cryoprotective solution, and stored at −20 °C until histological processing. Coronal sections corresponding to interaural levels 19.65-14.25 mm (Paxinos macaque atlas, 2012) were selected, encompassing the region targeted by the injections.

Sections were acclimated to room temperature, rinsed in PBS1X, and incubated for 1 h in blocking solution containing 3% normal serum and 0.3% Triton X-100 (NGS for Monkey S, NDS for Monkey T). Sections were then incubated overnight with primary antibodies diluted in PBS1X containing 1% matching serum and 0.03% Triton X-100: rabbit anti-oxytocin (Immunostar, #20068; 1:1000) and mouse anti-HA (Cell Signaling, #2367; 1:100) for Monkey S; guinea pig anti-oxytocin (Synaptic Systems, #408004; 1:500) and rabbit anti-HA (Cell Signaling, #3724; 1:100) for Monkey T. The next day, sections were rinsed in PBS1X and incubated for 2 h at room temperature with secondary antibodies diluted in PBS containing 3% matching serum: goat anti-rabbit Alexa 488 (Abcam, 1:500) and goat anti-mouse Alexa 647 (Abcam, 1:500) for Monkey S; Cy3 donkey anti-rabbit (Jackson ImmunoResearch, 1:100) and Cy5 donkey anti-guinea pig (Jackson ImmunoResearch, 1:100) for Monkey T. Sections were finally rinsed and mounted with Fluoromount DAPI mounting medium. Fluorescent labeling was imaged with an LSM 700 confocal microscope and analyzed with Fiji.

### Behavioral task

#### Digit-tracking method

Digit-tracking was implemented as a touchscreen-based method for measuring visual exploration, in which stimuli were globally blurred, so that exploration required active sampling of local image information, providing a functional analogue of foveated visual exploration. Finger contact revealed a circular Gaussian window, centered slightly above the contact point to prevent occlusion, with maximal resolution at its center and progressively reduced resolution toward its boundary. The window position was updated continuously as the finger slid across the display, and the coordinates of its center were recorded as a proxy for gaze exploration. The implementation followed the framework of Lio et al.^55^ and a macaque-specific adaptation^54^, which demonstrated digit-tracking as an unconstrained and minimally invasive alternative to eye-tracking for reliably capturing image exploration. All parameter settings were identical to those described in Yang et al.^54^

#### Apparatus and procedure

The experimental setup was identical to that described in Yang et al.^54^ Briefly, stimuli were displayed on a Dell Latitude 7210 tablet (60 Hz refresh rate; 12.3″ diagonal; 1920 × 1280 pixels) integrated into a battery-powered mobile device equipped with a seating platform and a liquid-delivery system. The mobile device was positioned against the front of the home cage to give subjects direct access to the touchscreen.

Each trial began with the appearance of a green virtual button at the center of the screen on a black background. Touching the button initiated the presentation of the blurred stimulus image. The first contact with the image started a 4 s free exploration window, during which subjects were free to explore the stimulus using the digit-tracking method. At the end of this period, the image was removed and a drop of water was delivered. A fixed 2 s interval followed to allow reward consumption before the next trial began.

#### Stimuli and presentation

A total of 300 color images were used as stimuli, divided into three categories: face images (n = 210), animal images (n = 45), and macaque social scenes (n = 45). The face set consisted of 21 individual models: 13 macaques from the facility colony and 8 humans. Each model was photographed 10 times in a frontal view: 5 with an averted gaze and 5 with a direct gaze, depicting individuals of both sexes (male/female) and varying familiarity (familiar/unfamiliar). All face images were cropped to include the entire head, resized into a predefined frame using GIMP (GNU Image Manipulation Program), and presented on an eigengrau (dark gray) background to ensure consistent facial feature size and a uniform background across stimuli. The image set was intentionally diverse to maintain the curiosity and engagement of the subjects. The stimulus set was divided into two complementary subsets of 150 images. Face stimuli were distributed to ensure a balanced representation of species, gaze direction, sex, and familiarity within each subset. This design allowed each subset to serve independently as a balanced collection. During a session, images from one subset were presented in a pseudo-random order until completion. To maximize on-screen face size, images were resized at presentation to fit the screen dimensions, such that the face covered approximately 52° of vertical visual angle (16 cm height at a minimal eye–screen distance of 16.5 cm). For face stimuli, the display position (left, center, or right) was randomized across trials to prevent the emergence of a central bias and to encourage exploration across the screen. Thus, a session included 105 face images, while the remainder consisted of animal and social-scene images. Although all categories were displayed, analyses reported in this study focused exclusively on the exploration of face images.

#### Training

Prior to data collection, monkeys were trained to perform the digit-tracking task using an independent set of images. The duration of the free-exploration window was gradually increased over successive training sessions, beginning at 1 s and incremented in 1 s steps until the final duration of 4 s was reached. The exploration window was increased once subjects reliably completed 150 trials within a session. Training continued until each monkey was able to complete full sessions.

#### Experimental conditions and sessions

Following training, experimental sessions were conducted under pharmacological manipulations. Conditions were defined along two dimensions: pharmacological state (vehicle vs DREADD ligand) and phase relative to viral vector injection (pre- vs post-injection). Post-injection sessions began at least six weeks after viral vector delivery. A minimum of 315 face trials was collected for each pharmacological state-by-phase condition. A complete session contained 105 face trials, although data from incomplete sessions were retained. The two face image subsets were counterbalanced across conditions within each phase. The acquisition criterion was chosen to limit habituation and maintain engagement with the task. Monkeys S and T reached this criterion within three full sessions per condition, yielding 315 trials per condition. By contrast, Monkey Y did not consistently complete full post-operative sessions, requiring additional sessions to reach the target number of trials, yielding 532 vehicle trials and 384 DCZ trials.

### Data analysis and statistics

All analyses were conducted using MATLAB (R2022a, MathWorks) and R (version 4.5.1, R Core Team) using custom scripts.

#### Exploration maps

Exploration maps were generated from digit-tracking data on a trial-by-trial basis. The recorded aperture-center coordinates were first transformed from screen space to original image space. For each trial, the time spent at each image coordinate was accumulated and then smoothed with a Gaussian kernel (σ = 2.6°), identical to the kernel applied to define the foveal aperture during the task. This procedure generated a single-trial exploration map of smoothed time spent across the image. These maps were used for spatial analyses, while temporal analyses of transitions between regions of interest were performed directly on the time series of aperture coordinates.

#### Cliff’s delta maps

Spatial treatment effects on exploration maps were first quantified using Cliff’s delta (δ), a non-parametric effect size computed at each pixel by pairwise comparison of ligand and vehicle trials. For every possible pair, a score of +1 was assigned if the ligand value exceeded the vehicle value, −1 if smaller, and 0 if equal. δ was defined as the average of these comparisons:

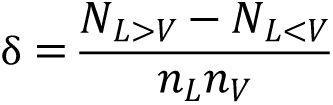

Here, N_L>V_ and N_L<V_ denote the numbers of trial pairs in which the ligand value was higher or lower than the vehicle value, respectively, and n_L_ and n_V_ denote the numbers of trials in each condition. δ ranges from −1 to +1, with δ = +1 indicating all ligand trials exceeded vehicle trials, δ = −1 the reverse, and δ = 0 no systematic difference. This measure captures how consistently exploration differed between treatment conditions across trials, irrespective of the magnitude of the difference. δ maps were lightly smoothed with a Gaussian filter for visualization. To visualize the most consistent treatment effects, regions corresponding to the highest and lowest 2.5% of δ values were displayed as connected clusters, separately for positive and negative values.

#### Spatially resolved within-subject treatment decoding

Treatment discriminability from exploration maps was evaluated using logistic classifiers. Exploration maps were downsampled to a 100 × 100 grid, transformed with a log1p non-linearity (x → log(1+x)) to reduce skewness, vectorized, and projected into a principal component analysis (PCA) feature space derived from the training data. Logistic classifiers were trained on PCA scores and evaluated using 10-fold cross-validation repeated 100 times. For each fold, performance was quantified by the receiver operating characteristic (ROC) curve and its area under the curve (AUC). ROC curves and AUC values were averaged across folds and repeats, and bootstrap confidence intervals were computed. Statistical significance was assessed using a permutation test in which training labels were shuffled and the complete shuffle-retrain-test procedure repeated 1000 times. To identify the spatial features contributing to treatment classification, logistic weights were back-projected from PCA space into the original exploration map space. Positive weights indicated contributions toward ligand classification, and negative weights toward vehicle classification. Spatial correspondence between decoder weights and Cliff’s delta maps was assessed using Pearson correlation.

#### Spatially resolved cross-subject treatment decoding

To assess generalization of treatment-related exploration patterns across individuals, logistic classifiers were trained on exploration maps from one subject and tested on another. Decoding was performed in both directions (subject A → subject B and subject B → subject A) and results were pooled across directions. Preprocessing followed the same procedure as the within-subject analysis (downsampling, log1p transform, PCA projection). Performance was quantified by ROC/AUC with bootstrap confidence intervals, and significance was tested against 1000 training label permutations per direction. To identify the spatial features supporting cross-subject generalization, an occlusion analysis was performed. For each decoding direction, small regions of the test maps were sequentially masked, and the resulting decrease in classifier performance (ΔAUC = AUCbaseline − AUCoccluded) was used to estimate the contribution of each location to decoding accuracy. Contribution maps were averaged across decoding directions. For visualization, ΔAUC was multiplied by 100 and expressed in percentage points.

#### Regions of interest (ROIs)

ROIs were defined a priori to cover the main facial features with equal size: eyes, nose, and mouth. An additional non-face ROI encompassed the remaining image area, including head and background. Spatial treatment-effect maps suggested a lateralized pattern of eye exploration. To capture this asymmetry, the original eyes ROI was post hoc divided into two equal regions (left and right eye). Lateralization analyses were conducted to examine overall exploration asymmetry. ROIs are illustrated on an example face image in Figure 2C.

#### ROI-based exploration analyses

Exploration time in each ROI was expressed as the proportion of the 4 s window spent in that region (bounded between 0 and 1). Because many trials contained zeros (no exploration in a given ROI), data were analyzed with a zero-inflated beta regression, a hurdle-type model that explicitly accounts for both zeros and continuous proportions in [0,1]. Separate models were fitted per subject for the available phases (before and/or after viral vector injection). The model included ROI, treatment, and their interaction as main factors, together with stimulus-related covariates (species, familiarity, gaze, sex, image position), a random intercept for session, and a random intercept for trial to account for clustering of ROI values within trials. Model formula:

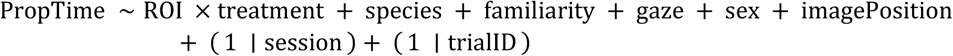

The regression included a conditional model for proportions (logit link) and a zero-component modeling the probability of no exploration, with dispersion estimated separately for each ROI. The main analyses targeted the ROI × treatment interaction, while exploratory three-way interactions with stimulus covariates (treatment × ROI × covariate) were tested individually but none were significant. Model adequacy was checked through residual diagnostics, convergence assessments, collinearity checks, and comparison with simpler alternatives. The model was decomposed into three complementary components: the probability that an ROI was explored (hit probability), the proportion of time spent in an ROI conditional on exploration, and the overall proportion of time (including zeros). Although all three components were estimated, analyses focused on hit probability and conditional exploration time, which provide complementary information about exploration occurrence and duration. Within each component, marginal means and post-hoc contrasts were computed for all ROIs, comparing ligand against vehicle. Effects were expressed as relative change (ligand/vehicle − 1), such that 0 indicated no effect. P-values were adjusted using the Benjamini–Hochberg procedure. Implementation used the R packages glmmTMB for model fitting, marginaleffects for marginal means and contrasts, and DHARMa for diagnostics.

#### ROI-based transition analyses

Transition analyses relied on within-trial aperture time series rather than spatial maps. Each time point was assigned to an ROI; contiguous periods within the same ROI were collapsed; a transition was defined at each change of ROI. Transitions were modeled as first-order choices, predicting the next ROI from the current ROI. Separate models were fitted per subject for the available phases (before and/or after viral vector injection). A multinomial logistic regression was used with current ROI, treatment, and their interaction as main factors, together with stimulus-related covariates (species, familiarity, gaze, sex, image position) and a quadratic term for within-trial step. Model formula:

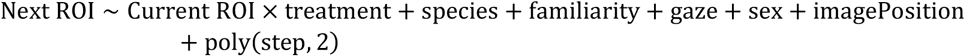

Trial-level random effects were not identifiable in the multinomial framework and were therefore omitted. A session random intercept was explored but excluded because sparse transition counts and quasi-complete separation prevented model convergence. Model adequacy was checked with standard diagnostics and comparisons to simpler specifications. Predicted transition probabilities were obtained for every current-to-next-ROI pair under each treatment. For each pair, marginal means and post-hoc contrasts tested ligand versus vehicle as differences in probability (ΔP; 0 indicates no effect), with Benjamini–Hochberg correction. To summarize how transitions were distributed across ROIs, in-strength was computed from treatment-specific transition matrices as the proportion of incoming transitions landing in each ROI. Ligand vs vehicle differences were assessed with permutation tests (1000 treatment-label shuffles) and Benjamini–Hochberg correction across ROIs. Implementation used the R packages nnet for model fitting and emmeans for marginal means and contrasts.

#### ROI-based cross-subject treatment decoding

To assess whether treatment effects generalized across subjects at the level of facial features, logistic classifiers were trained on ROI-based measures from one subject and tested on another. Decoding was performed in both directions, and results were pooled across directions. Features included exploration time, hit probability, ROI in-strength and transition probabilities. Statistical significance was assessed using the same permutation procedure as for spatial cross-subject decoding. To identify the ROIs contributing most strongly to cross-subject generalization, decoding performance was evaluated using individual ROIs and by measuring the decrease in performance associated with their removal. Results were averaged across directions. AUC values below 0.5 indicate systematic reversal of the decoded pattern between training and test datasets.

#### Lateralization index (LI) analyses

Two time-based lateralization indices were computed per trial: a screen-referenced LI (relative to the screen midline) and a face-referenced LI (relative to the face midline on screen, irrespective of screen position). LI was computed from exploration time as:

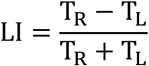

Here, T_R_ and T_L_ denote the exploration times on the right and left sides of the relevant midline, respectively. LI values range from −1 to 1, with negative and positive values indicating leftward and rightward biases, respectively. For each LI (screen- and face-referenced), mean LI values for ligand and vehicle were estimated with bootstrap confidence intervals and ligand-vehicle differences were assessed with stratified permutation tests by image position (1000 permutations). P-values were adjusted using the Benjamini–Hochberg procedure.

## DATA AND CODE AVAILABILITY

- All data reported in this paper will be shared by the corresponding author upon request.
- All code is available from the corresponding author upon request.
- Any additional information required to reanalyze the data reported in this paper is available from the corresponding author upon request.

## ACKNOWLEDGMENTS

The authors thank M.-V. Baussant, S. Duray and C. Ruppé for administrative support; and F. Francioli, L. Cabrol, M. Robiani and M.-L. Rouge for assistance with animal welfare and care; F. Lavenne and L. Tremblay for technical help. This work was supported by grants from the European Research Council (ERC) under the European Union’s Horizon 2020 research and innovation programme (grant agreement no. 885746 to J.-R.D.) and Horizon Europe research and innovation programme (grant agreement no. 101071777 to V.G. and A.S.).

## AUTHOR CONTRIBUTIONS

Conceptualization, A.A., A.S. and J.-R.D.; methodology, A.A., E.D., H.E., C.D., V.G. and J.-R.D.; investigation, A.A., E.D., H.E. and C.D.; formal analysis, A.A.; visualization, A.A., H.E.; writing—original draft, A.A.; writing—review & editing, A.A., E.D., H.E., C.D., V.G., A.S., and J.-R.D.; funding acquisition, V.G., A.S. and J.-R.D.; resources, V.G., A.S. and J.-R.D.; supervision, A.S. and J.-R.D.

## SUPPLEMENTARY INFORMATION

**Supplementary Figure 1.**
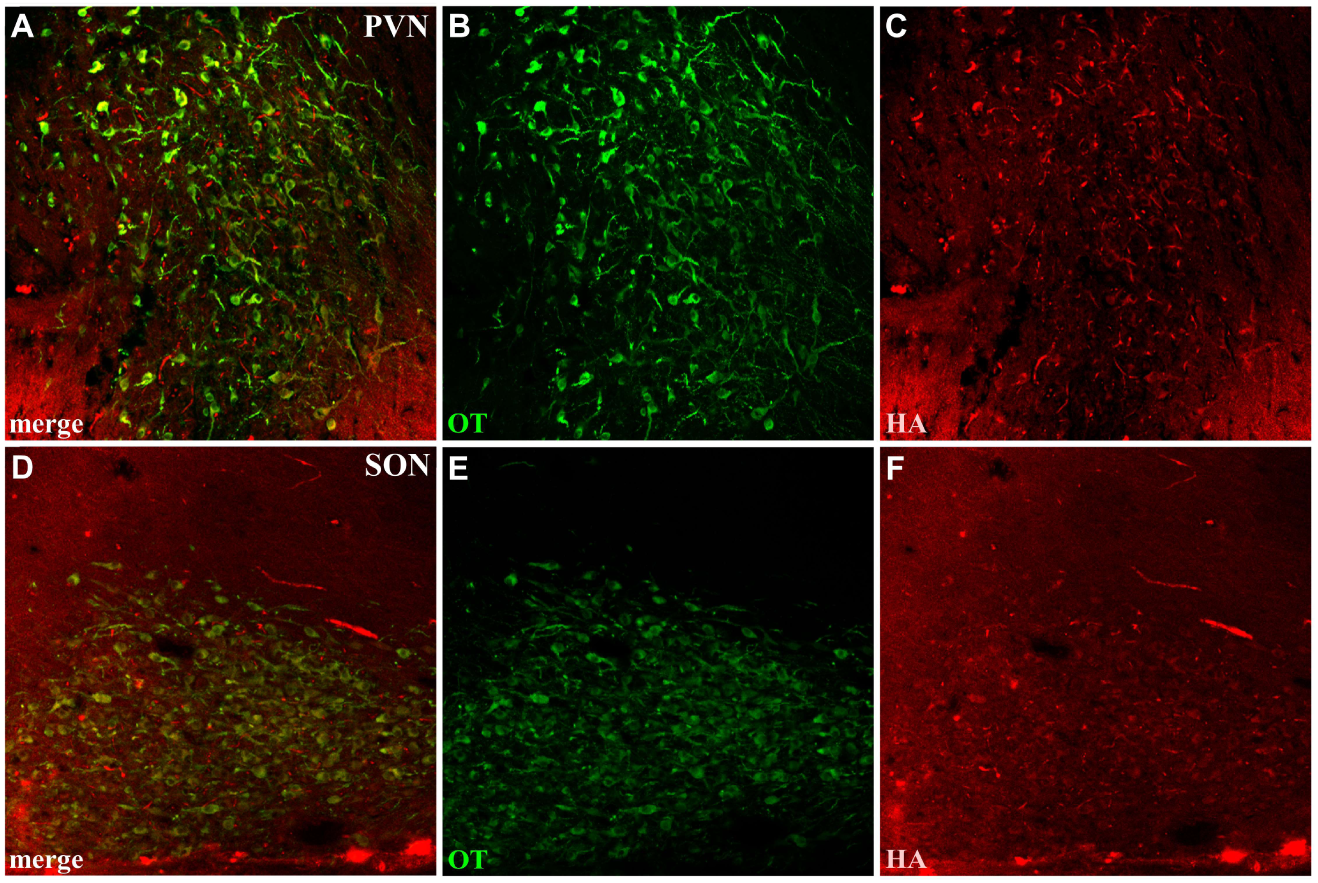
Double immunofluorescence labeling of OT (green) and HA-tag (red) in Monkey S. **A–C.** Images showing OT and HA-tag labeling in the PVN, shown as a merged image in A and as single-channel images in B and C. **D–F.** Images showing OT and HA-tag labeling in the SON, shown as a merged image in D and as single-channel images in E and F. Images were acquired at an approximate Paxinos interaural level of +17.45 mm.

**Supplementary Figure 2.**
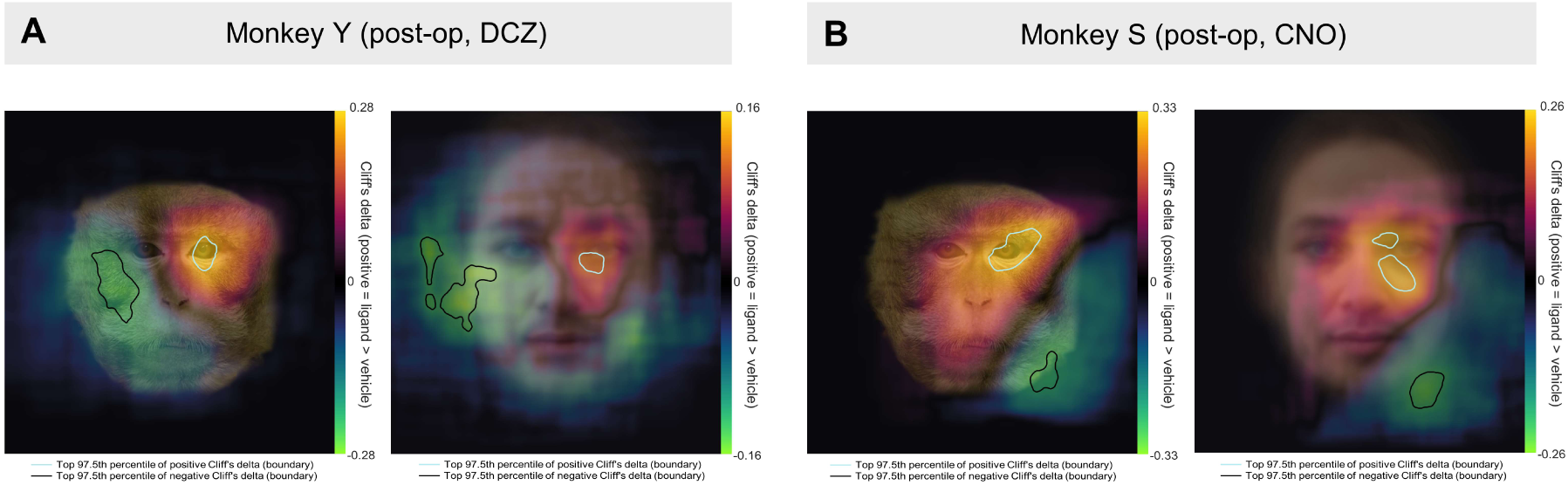
OT spatial signatures on macaque and human faces. **A, B.** Panels A and B correspond to Monkey Y and Monkey S. For each monkey, the left and right plots show Cliff’s delta (δ) maps for macaque and human faces, respectively, overlaid on representative face images. For visualization, the human image was a computer-generated average of the face stimuli used in the task and does not depict an identifiable individual. δ values, computed at each pixel across all ligand–vehicle trial pairs, reflect the direction and consistency of treatment-related differences. Both monkeys were injected with AAV9 carrying the OTpr-HA-hM3Dq construct in the PVN-SON region. Monkey Y was tested with DCZ (0.1 mg/kg) and Monkey S with CNO (3 mg/kg) as DREADD ligands.

**Supplementary Figure 3.**
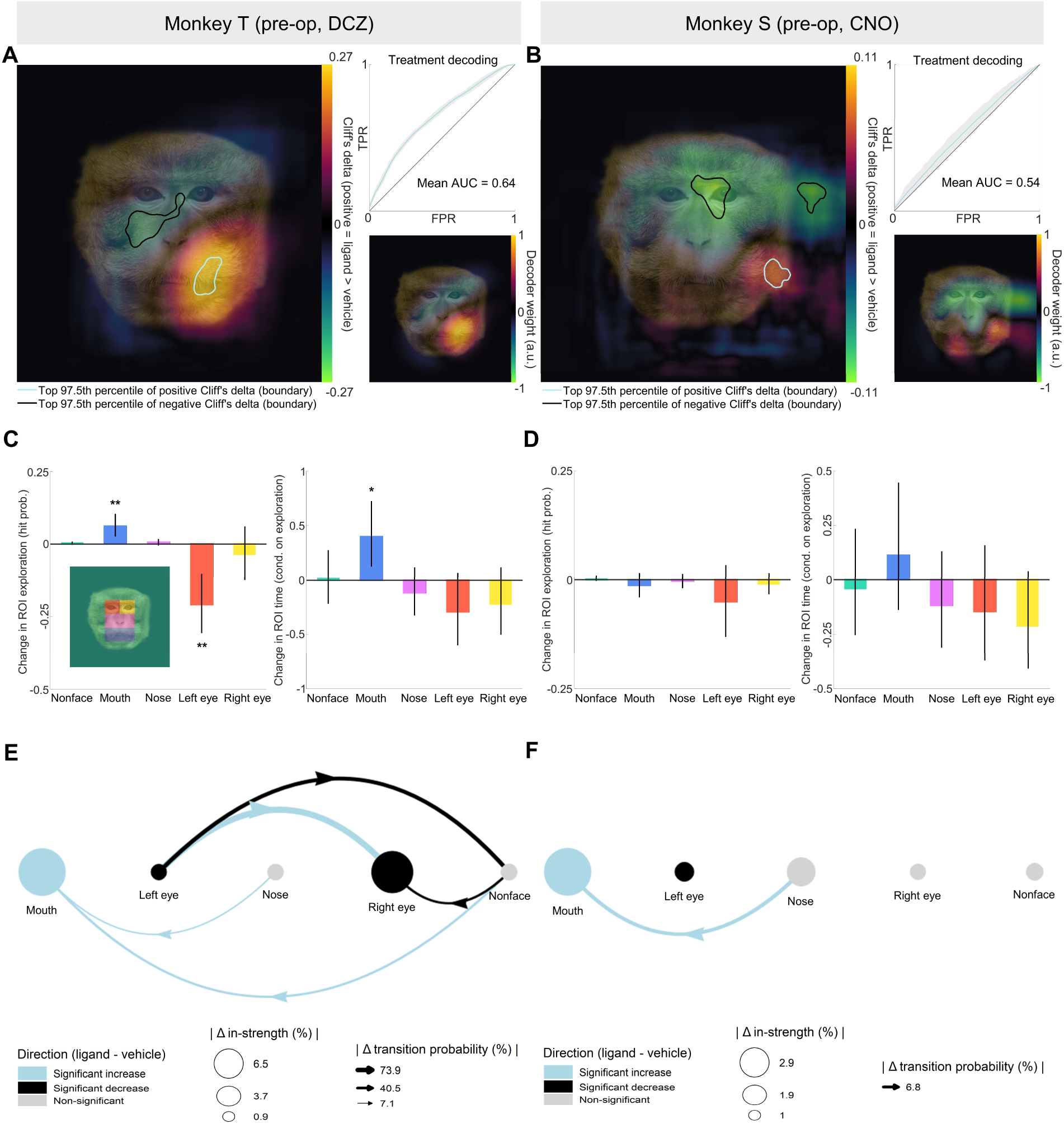
Pre-operative effects of DREADD ligands on face exploration before viral vector delivery. **A, B.** Spatially resolved analysis of exploration maps in Monkey T (DCZ) and Monkey S (CNO) before AAV vector delivery. Left: Cliff’s delta (δ) map showing non-parametric effect sizes computed at each pixel across all ligand–vehicle trial pairs, reflecting the direction and consistency of treatment-related differences. Top right: ROC curves of logistic classifiers trained to decode treatment condition from single-trial exploration maps (gray shading: confidence intervals). Bottom right: Decoder weight maps highlighting the spatial features contributing most strongly to classification. In Monkey T, δ maps and decoder weights emphasized the lower face and mouth region, while negative values overlapped the eyes and right side of the nose, indicating reduced eye-directed exploration under DCZ. Decoding significantly discriminated ligand from vehicle trials (AUC = 0.64, p = 0.004, permutation test), and decoder weights strongly correlated with δ maps (r = 0.96, p < 0.001). In Monkey S, δ values were small in magnitude (absolute maximum = 0.11), with weak negative values over the eyes and nose and only limited increases on the right side of the head. Decoding performance was weak but significant (AUC = 0.54, p = 0.046). **C, D.** ROI exploration metrics estimated with a hurdle model (zero-inflated beta regression). Left: relative change (ligand / vehicle − 1) in the probability that each ROI was explored at least once during the 4 s exploration window (hit probability). Right: relative change in the proportion of exploration time spent within each ROI when explored (conditional proportion of time). Vertical lines indicate 95% confidence intervals; asterisks denote significance after Benjamini–Hochberg correction (*p < 0.05, **p < 0.01, ***p < 0.001). In Monkey T, DCZ reduced visits to the eyes, particularly the left eye, and increased both the probability and duration of mouth exploration. In Monkey S, CNO induced no reliable changes in eye exploration or exploration time allocation. **E, F.** ROI in-strength and transition analyses. Nodes represent ROI in-strength (proportion of incoming transitions landing in each ROI), and edges represent transition probabilities between ROIs. Only significant transitions after Benjamini–Hochberg correction are shown. In Monkey T, DCZ increased exploration flows toward the mouth, notably from the nose and non-face regions, while reducing transitions toward the eyes. In Monkey S, only minor transition changes were observed, including a small increase in transitions toward the mouth region. Together, these pre-operative analyses show that ligand administration alone did not reproduce the post-operative enhancement of eye exploration observed after DREADD-mediated activation of OT neurons. When present, ligand effects were heterogeneous and predominantly involved increased exploration of lower-face regions rather than the eyes.

**Supplementary Figure 4.**
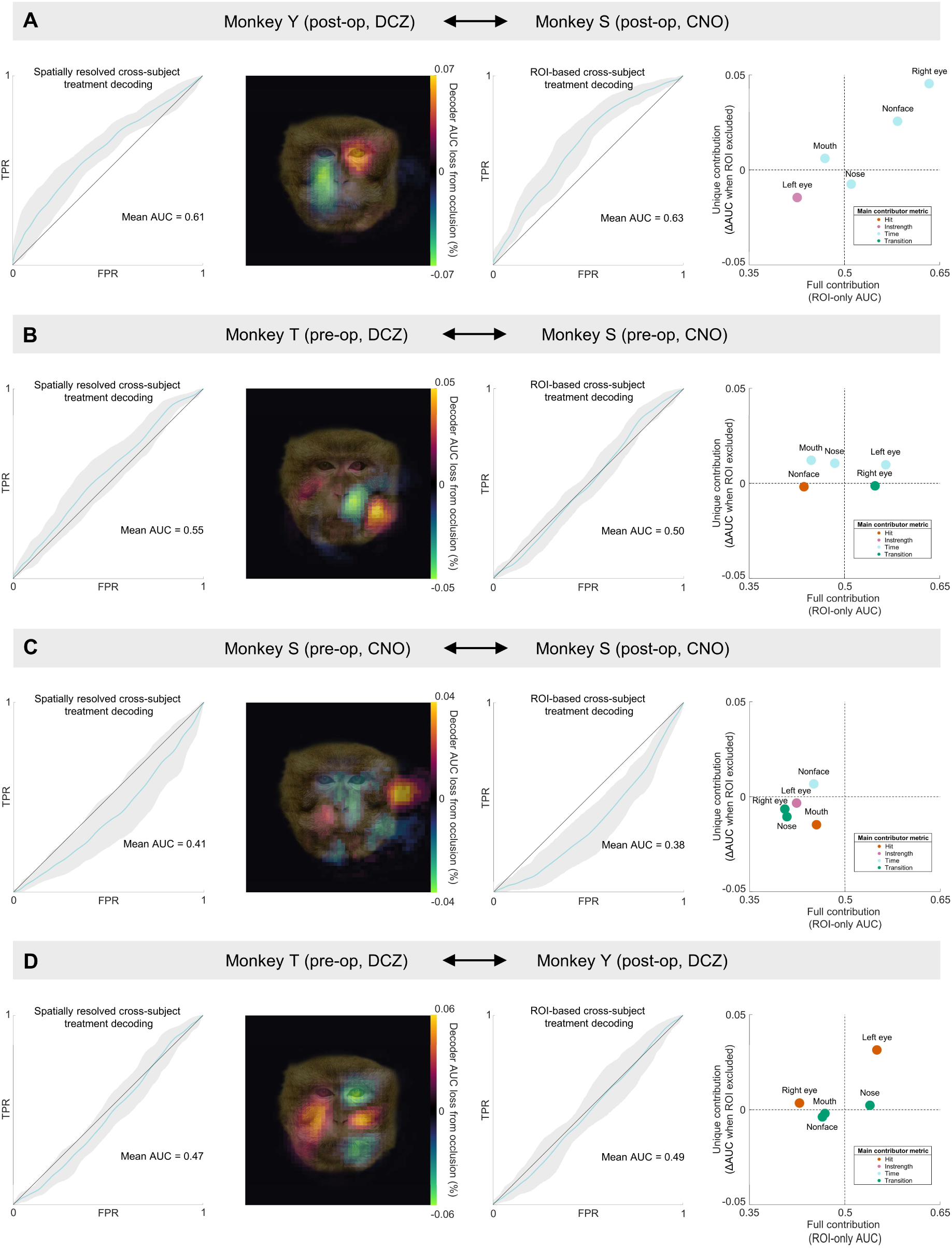
*G*eneralization of treatment effects. **A–D.** Each row shows one pairing, with subjects and operation status indicated in the row labels. Columns show four analyses. **First:** ROC curve for treatment decoding across the pairing with a logistic classifier using exploration maps. **Second:** Occlusion map showing the change in AUC (ΔAUC), expressed in percentage points (pp), when exploration at specific spatial locations is masked in the test set, identifying locations where treatment effects converge across the pairing (ΔAUC > 0, masking reduces generalization performance) or diverge (ΔAUC < 0, masking improves generalization performance). **Third:** ROC curve for treatment decoding across the pairing with a logistic classifier using ROI × metric features (hit, time, in-strength, and transition for each ROI). Fourth: ROI contribution plot. The x-axis represents ROI full contribution, defined as the AUC of a logistic classifier trained only on metrics from that ROI. The y-axis represents ROI unique contribution, defined as the change in AUC when the ROI is excluded from the full model trained on all ROI × metric features. Quadrants summarize treatment-effect patterns: top right, similar effects; top left, divergent effects with a consistent cross-ROI pattern; bottom right, consistent but non-unique effects; bottom left, opposite effects. For all decoding analyses, classifiers were trained on one member of each pairing and tested on the other, and vice versa. Gray shading indicates confidence intervals for ROC curves.

## REFERENCES

1. Silk, J.B. (2007). The adaptive value of sociality in mammalian groups. Phil. Trans. R. Soc. B 362, 539–559. 10.1098/rstb.2006.1994.

2. Ghazanfar, A.A., and Santos, L.R. (2004). Primate brains in the wild: the sensory bases for social interactions. Nat Rev Neurosci 5, 603–616. 10.1038/nrn1473.

3. Emery, N.J. (2000). The eyes have it: the neuroethology, function and evolution of social gaze. Neuroscience & Biobehavioral Reviews 24, 581–604. 10.1016/S0149-7634(00)00025-7.

4. Itier, R.J., and Batty, M. (2009). Neural bases of eye and gaze processing: The core of social cognition. Neuroscience & Biobehavioral Reviews 33, 843–863. 10.1016/j.neubiorev.2009.02.004.

5. Dunbar, R. I. M., & Shultz, S. (2007). Evolution in the social brain. Science, 317, 1344–1347. 10.1126/science.1145463

6. Deen, B., Schwiedrzik, C.M., Sliwa, J., and Freiwald, W.A. (2023). Specialized Networks for Social Cognition in the Primate Brain. Annu. Rev. Neurosci. 46, 381–401. 10.1146/annurev-neuro-102522-121410.

7. Freeman, S.M., Inoue, K., Smith, A.L., Goodman, M.M., and Young, L.J. (2014). The neuroanatomical distribution of oxytocin receptor binding and mRNA in the male rhesus macaque (Macaca mulatta). Psychoneuroendocrinology 45, 128–141. 10.1016/j.psyneuen.2014.03.023.

8. Quintana, D.S., Rokicki, J., van der Meer, D., Alnæs, D., Kaufmann, T., Córdova-Palomera, A., Dieset, I., Andreassen, O.A., and Westlye, L.T. (2019). Oxytocin pathway gene networks in the human brain. Nature Communications 10. 10.1038/s41467-019-08503-8.

9. Son, S., Manjila, S.B., Newmaster, K.T., Wu, Y., Vanselow, D.J., Ciarletta, M., Anthony, T.E., Cheng, K.C., and Kim, Y. (2022). Whole-Brain Wiring Diagram of Oxytocin System in Adult Mice. J. Neurosci. 42, 5021–5033. 10.1523/JNEUROSCI.0307-22.2022.

10. Caffé, A. R., Van Ryen, P. C., Van der Woude, T. P., and van Leeuwen, F. W. (1989). Vasopressin and oxytocin systems in the brain and upper spinal cord of *Macaca fascicularis*. J. Comp. Neurol. 287, 302–325. 10.1002/cne.902870304

11. Rogers, C.N., Ross, A.P., Sahu, S.P., Siegel, E.R., Dooyema, J.M., Cree, M.A., Stopa, E.G., Young, L.J., Rilling, J.K., Albers, H.E., et al. (2018). Oxytocin- and arginine vasopressin-containing fibers in the cortex of humans, chimpanzees, and rhesus macaques. American Journal of Primatology 80. 10.1002/ajp.22875.

12. Lefevre, A., Meza, J., and Miller, C.T. (2024). Long-range projections of oxytocin neurons in the marmoset brain. J Neuroendocrinology 36, e13397. 10.1111/jne.13397.

13. Froemke, R.C., and Young, L.J. (2021). Oxytocin, Neural Plasticity, and Social Behavior. Annu. Rev. Neurosci. 44, 359–381. 10.1146/annurev-neuro-102320-102847.

14. Quintana, D.S., and Guastella, A.J. (2020). An Allostatic Theory of Oxytocin. Trends in Cognitive Sciences. 10.1016/j.tics.2020.03.008.

15. Menon, R., and Neumann, I.D. (2023). Detection, processing and reinforcement of social cues: regulation by the oxytocin system. Nat. Rev. Neurosci. 10.1038/s41583-023-00759-w.

16. Stoop, R. (2012). Neuromodulation by Oxytocin and Vasopressin. Neuron 76, 142–159. 10.1016/j.neuron.2012.09.025.

17. Krabichler Q, Grinevich V 2025. Evolution of Neuropeptide Signaling: From a Single Cell to Mammals. In: Grinevich V, Oliveira R editors. Evolutionary and Comparative Neuroendocrinology. Masterclass in Neuroendocrinology, volume 17. Springer, Cham. pp 3 to 43. 10.1007/978-3-031-80209-6_1

18. Soumier, A., Habart, M., Lio, G., Demily, C., and Sirigu, A. (2022). Differential fate between oxytocin and vasopressin cells in the developing mouse brain. iScience 25, 103655. 10.1016/j.isci.2021.103655.

19. Tang, Y., Benusiglio, D., Lefevre, A., Hilfiger, L., Althammer, F., Bludau, A., Hagiwara, D., Baudon, A., Darbon, P., Schimmer, J., et al. (2020). Social touch promotes interfemale communication via activation of parvocellular oxytocin neurons. Nature Neuroscience. 10.1038/s41593-020-0674-y.

20. Duque-Wilckens, N., Torres, L.Y., Yokoyama, S., Minie, V.A., Tran, A.M., Petkova, S.P., Hao, R., Ramos-Maciel, S., Rios, R.A., Jackson, K., et al. (2020). Extrahypothalamic oxytocin neurons drive stress-induced social vigilance and avoidance. Proc. Natl. Acad. Sci. U.S.A. 117, 26406–26413. 10.1073/pnas.2011890117.

21. Osakada, T., Yan, R., Jiang, Y., Wei, D., Tabuchi, R., Dai, B., Wang, X., Zhao, G., Wang, C.X., Liu, J.-J., et al. (2024). A dedicated hypothalamic oxytocin circuit controls aversive social learning. Nature. 10.1038/s41586-023-06958-w.

22. Kendrick, K.M., Guastella, A.J., and Becker, B. (2017). Overview of Human Oxytocin Research. In Behavioral Pharmacology of Neuropeptides: Oxytocin Current Topics in Behavioral Neurosciences., R. Hurlemann and V. Grinevich, eds. (Springer International Publishing), pp. 321–348. 10.1007/7854_2017_19.

23. Leng, G., and Ludwig, M. (2016). Intranasal Oxytocin: Myths and Delusions. Biological Psychiatry 79, 243–250. 10.1016/j.biopsych.2015.05.003.

24. Walum, H., Waldman, I.D., and Young, L.J. (2016). Statistical and Methodological Considerations for the Interpretation of Intranasal Oxytocin Studies. Biological Psychiatry 79, 251–257. 10.1016/j.biopsych.2015.06.016.

25. Insel, T.R. (2010). The Challenge of Translation in Social Neuroscience: A Review of Oxytocin, Vasopressin, and Affiliative Behavior. Neuron. 10.1016/j.neuron.2010.03.005.

26. Freeman, S.M., and Young, L.J. (2016). Comparative Perspectives on Oxytocin and Vasopressin Receptor Research in Rodents and Primates: Translational Implications. Journal of Neuroendocrinology 28. 10.1111/jne.12382.

27. Grinevich, V., and Neumann, I.D. (2021). Brain oxytocin: how puzzle stones from animal studies translate into psychiatry. Mol Psychiatry 26, 265–279. 10.1038/s41380-020-0802-9.

28. Phillips, K.A., Bales, K.L., Capitanio, J.P., Conley, A., Czoty, P.W., ‘T Hart, B.A., Hopkins, W.D., Hu, S., Miller, L.A., Nader, M.A., et al. (2014). Why primate models matter. American J Primatol 76, 801–827. 10.1002/ajp.22281.

29. Putnam, P.T., Young, L.J., and Gothard, K.M. (2018). Bridging the gap between rodents and humans: The role of non-human primates in oxytocin research. Am J Primatol 80, e22756. 10.1002/ajp.22756.

30. Chang, S.W.C., and Platt, M.L. (2014). Oxytocin and social cognition in rhesus macaques: Implications for understanding and treating human psychopathology. Brain Research 1580, 57–68. 10.1016/j.brainres.2013.11.006.

31. Tremblay, S., Acker, L., Afraz, A., Albaugh, D.L., Amita, H., Andrei, A.R., Angelucci, A., Aschner, A., Balan, P.F., Basso, M.A., et al. (2020). An Open Resource for Non-human Primate Optogenetics. Neuron 108, 1075–1090.e6. 10.1016/j.neuron.2020.09.027.

32. Raper, J., and Galvan, A. (2022). Applications of chemogenetics in non-human primates. Current Opinion in Pharmacology 64, 102204. 10.1016/j.coph.2022.102204.

33. Knobloch, H.S., Charlet, A., Hoffmann, L.C., Eliava, M., Khrulev, S., Cetin, A.H., Osten, P., Schwarz, M.K., Seeburg, P.H., Stoop, R., et al. (2012). Evoked axonal oxytocin release in the central amygdala attenuates fear response. Neuron 73, 553–566. 10.1016/j.neuron.2011.11.030.

34. Eliava, M., Melchior, M., Knobloch-Bollmann, H.S., Wahis, J., da Silva Gouveia, M., Tang, Y., Ciobanu, A.C., Triana del Rio, R., Roth, L.C., Althammer, F., et al. (2016). A New Population of Parvocellular Oxytocin Neurons Controlling Magnocellular Neuron Activity and Inflammatory Pain Processing. Neuron 89, 1291–1304. 10.1016/j.neuron.2016.01.041.

35. Hasan, M.T., Althammer, F., Silva da Gouveia, M., Goyon, S., Eliava, M., Lefevre, A., Kerspern, D., Schimmer, J., Raftogianni, A., Wahis, J., et al. (2019). A Fear Memory Engram and Its Plasticity in the Hypothalamic Oxytocin System. Neuron 103, 133–146.e8. 10.1016/j.neuron.2019.04.029.

36. Grund, T., Tang, Y., Benusiglio, D., Althammer, F., Probst, S., Oppenländer, L., Neumann, I.D., and Grinevich, V. (2019). Chemogenetic activation of oxytocin neurons: Temporal dynamics, hormonal release, and behavioral consequences. Psychoneuroendocrinology 106, 77–84. 10.1016/j.psyneuen.2019.03.019.

37. Nishimura, H., Yoshimura, M., Shimizu, M., Sanada, K., Sonoda, S., Nishimura, K., Baba, K., Ikeda, N., Motojima, Y., Maruyama, T., et al. (2022). Endogenous oxytocin exerts anti-nociceptive and anti-inflammatory effects in rats. Commun Biol 5, 907. 10.1038/s42003-022-03879-8.

38. Guastella, A.J., Mitchell, P.B., and Dadds, M.R. (2008). Oxytocin Increases Gaze to the Eye Region of Human Faces. Biological Psychiatry 63, 3–5. 10.1016/j.biopsych.2007.06.026.

39. Gamer, M., Zurowski, B., and Büchel, C. (2010). Different amygdala subregions mediate valence-related and attentional effects of oxytocin in humans. Proc. Natl. Acad. Sci. U.S.A. 107, 9400–9405. 10.1073/pnas.1000985107.

40. Andari E, Duhamel JR, Zalla T, Herbrecht E, Leboyer M, and Sirigu A. (2010). Promoting social behavior with oxytocin in high-functioning autism spectrum disorders. Proceedings of the National Academy of Sciences of the United States of America 107, 4389–4394. 10.1073/pnas.0910249107

41. Chang, S.W.C., Barter, J.W., Becket Ebitz, R., Watson, K.K., and Platt, M.L. (2012). Inhaled oxytocin amplifies both vicarious reinforcement and self reinforcement in rhesus macaques (Macaca mulatta). Proceedings of the National Academy of Sciences of the United States of America 109, 959–964. 10.1073/pnas.1114621109.

42. Dal Monte, O., Noble, P.L., Costa, V.D., and Averbeck, B.B. (2014). Oxytocin enhances attention to the eye region in rhesus monkeys. Frontiers in Neuroscience. 10.3389/fnins.2014.00041.

43. Kotani, M., Shimono, K., Yoneyama, T., Nakako, T., Matsumoto, K., Ogi, Y., Konoike, N., Nakamura, K., and Ikeda, K. (2017). An eye tracking system for monitoring face scanning patterns reveals the enhancing effect of oxytocin on eye contact in common marmosets. Psychoneuroendocrinology 83, 42–48. 10.1016/j.psyneuen.2017.05.009.

44. Porffy, L.A., Bell, V., Coutrot, A., Wigton, R., D’Oliveira, T., Mareschal, I., and Shergill, S.S. (2020). In the eye of the beholder? Oxytocin effects on eye movements in schizophrenia. Schizophrenia Research 216, 279–287. 10.1016/j.schres.2019.11.044.

45. Brooks, J., Kano, F., Sato, Y., Yeow, H., Morimura, N., Nagasawa, M., Kikusui, T., and Yamamoto, S. (2021). Divergent effects of oxytocin on eye contact in bonobos and chimpanzees. Psychoneuroendocrinology 125, 105119. 10.1016/j.psyneuen.2020.105119.

46. Marsh, N., Scheele, D., Postin, D., Onken, M., and Hurlemann, R. (2021). Eye-Tracking Reveals a Role of Oxytocin in Attention Allocation Towards Familiar Faces. Front. Endocrinol. 12, 629760. 10.3389/fendo.2021.629760.

47. Dal Monte, O., Piva, M., Anderson, K.M., Tringides, M., Holmes, A.J., and Chang, S.W.C. (2017). Oxytocin under opioid antagonism leads to supralinear enhancement of social attention. Proceedings of the National Academy of Sciences of the United States of America 114, 5247– 5252. 10.1073/pnas.1702725114.

48. Parr, L.A., Brooks, J.M., Jonesteller, T., Moss, S., Jordano, J.O., and Heitz, T.R. (2016). Effects of chronic oxytocin on attention to dynamic facial expressions in infant macaques. Psychoneuroendocrinology 74, 149–157. 10.1016/j.psyneuen.2016.08.028.

49. Jiang, Y., and Platt, M.L. (2018). Oxytocin and vasopressin flatten dominance hierarchy and enhance behavioral synchrony in part via anterior cingulate cortex. Scientific Reports 8. 10.1038/s41598-018-25607-1.

50. Jiang, Y., and Platt, M.L. (2018). Oxytocin and vasopressin increase male-directed threats and vocalizations in female macaques. Scientific Reports 8, 1–14. 10.1038/s41598-018-36332-0.

51. Jiang, Y., Sheng, F., Belkaya, N., and Platt, M.L. (2022). Oxytocin and testosterone administration amplify viewing preferences for sexual images in male rhesus macaques. Phil. Trans. R. Soc. B 377, 20210133. 10.1098/rstb.2021.0133.

52. Jongerius, C., Hillen, M.A., Smets, E.M.A., Mol, M.J., Kooij, E.S., De Nood, M.A., Dalmaijer, E.S., Fliers, E., Romijn, J.A., and Quintana, D.S. (2023). Nasal oxytocin administration does not influence eye gaze or perceived relationship of male volunteers with physicians in a simulated online consultation: a randomized, placebo-controlled trial. Endocrine Connections, EC-22-0377. 10.1530/EC-22-0377.

53. Sosnowski, M.J., Kano, F., and Brosnan, S.F. (2022). Oxytocin and social gaze during a dominance categorization task in tufted capuchin monkeys. Front. Psychol. 13, 977771. 10.3389/fpsyg.2022.977771.

54. Yang, Y., Ameloot, A., Lio, G., Sirigu, A., and Duhamel, J.-R. (2026). Digit-tracking reveals curiosity-driven visual attention in macaque monkeys. Sci Rep. 10.1038/s41598-026-57654-4.

55. Lio, G., Fadda, R., Doneddu, G., Duhamel, J.R., and Sirigu, A. (2019). Digit-tracking as a new tactile interface for visual perception analysis. Nature Communications 10, 1–13. 10.1038/s41467-019-13285-0.

56. Nagai, Y., Miyakawa, N., Takuwa, H., Hori, Y., Oyama, K., Ji, B., Takahashi, M., Huang, X.P., Slocum, S.T., DiBerto, J.F., et al. (2020). Deschloroclozapine, a potent and selective chemogenetic actuator enables rapid neuronal and behavioral modulations in mice and monkeys. Nature Neuroscience 23, 1157–1167. 10.1038/s41593-020-0661-3.

57. Upright, N.A., and Baxter, M.G. (2020). Effect of chemogenetic actuator drugs on prefrontal cortex-dependent working memory in nonhuman primates. Neuropsychopharmacol. 45, 1793– 1798. 10.1038/s41386-020-0660-9.

58. Frischen, A., Bayliss, A.P., and Tipper, S.P. (2007). Gaze cueing of attention: Visual attention, social cognition, and individual differences. Psychological Bulletin 133, 694–724. 10.1037/0033-2909.133.4.694.

59. Bartz, J.A., Zaki, J., Bolger, N., and Ochsner, K.N. (2011). Social effects of oxytocin in humans: Context and person matter. Trends in Cognitive Sciences. 10.1016/j.tics.2011.05.002.

60. Grinevich, V., Desarménien, M.G., Chini, B., Tauber, M., and Muscatelli, F. (2015). Ontogenesis of oxytocin pathways in the mammalian brain: Late maturation and psychosocial disorders. Frontiers in Neuroanatomy 8, 1–17. 10.3389/fnana.2014.00164.

61. Costa, M., Gomez, A., Barat, E., Lio, G., Duhamel, J.-R., and Sirigu, A. (2018). Implicit preference for human trustworthy faces in macaque monkeys. Nat Commun 9, 4529. 10.1038/s41467-018-06987-4.

62. Howarth, E.R.I., Szott, I.D., Witham, C.L., Wilding, C.S., and Bethell, E.J. (2023). Genetic polymorphisms in the serotonin, dopamine and opioid pathways influence social attention in rhesus macaques (Macaca mulatta). PLoS ONE 18, e0288108. 10.1371/journal.pone.0288108.

63. Liu, S., Huang, J., Chen, S., Platt, M.L., and Yang, Y. (2025). Multi-dimensional social relationships shape social attention in monkeys. eLife 14, RP104460. 10.7554/eLife.104460.

64. Bethell, E.J., Cassidy, L.C., Brockhausen, R.R., and Pfefferle, D. (2019). Toward a Standardized Test of Fearful Temperament in Primates: A Sensitive Alternative to the Human Intruder Task for Laboratory-Housed Rhesus Macaques (Macaca mulatta). Front. Psychol. 10, 1051. 10.3389/fpsyg.2019.01051.

65. Kampe, K.K.W., Frith, C.D., Dolan, R.J., and Frith, U. (2001). Reward value of attractiveness and gaze. Nature 413, 589–589. 10.1038/35098149.

66. Scheele, D., Wille, A., Kendrick, K.M., Stoffel-Wagner, B., Becker, B., Güntürkün, O., Maier, W., and Hurlemann, R. (2013). Oxytocin enhances brain reward system responses in men viewing the face of their female partner. Proceedings of the National Academy of Sciences of the United States of America 110, 20308–20313. 10.1073/pnas.1314190110.

67. Ballesta, S., and Duhamel, J.R. (2015). Rudimentary empathy in macaques’ social decision-making. Proceedings of the National Academy of Sciences of the United States of America 112, 15516–15521. 10.1073/pnas.1504454112.

68. Hung, L.W., Neuner, S., Polepalli, J.S., Beier, K.T., Wright, M., Walsh, J.J., Lewis, E.M., Luo, L., Deisseroth, K., Dölen, G., et al. (2017). Gating of social reward by oxytocin in the ventral tegmental area. Science 357, 1406–1411. 10.1126/science.aan4994.

69. Nardou, R., Lewis, E.M., Rothhaas, R., Xu, R., Yang, A., Boyden, E., and Dölen, G. (2019). Oxytocin-dependent reopening of a social reward learning critical period with MDMA. Nature 569, 116–120. 10.1038/s41586-019-1075-9.

70. Lewis, E.M., Stein-O’Brien, G.L., Patino, A.V., Nardou, R., Grossman, C.D., Brown, M., Bangamwabo, B., Ndiaye, N., Giovinazzo, D., Dardani, I., et al. (2020). Parallel Social Information Processing Circuits Are Differentially Impacted in Autism. Neuron 108, 659–675.e6. 10.1016/j.neuron.2020.10.002.

71. Ellenbogen, M.A., Linnen, A.M., Grumet, R., Cardoso, C., and Joober, R. (2012). The acute effects of intranasal oxytocin on automatic and effortful attentional shifting to emotional faces. Psychophysiology 49, 128–137. 10.1111/j.1469-8986.2011.01278.x.

72. Ebitz, R.B., Watson, K.K., and Platt, M.L. (2013). Oxytocin blunts social vigilance in the rhesus macaque. Proc. Natl. Acad. Sci. U.S.A. 110, 11630–11635. 10.1073/pnas.1305230110.

73. Parr, L.A., Modi, M., Siebert, E., and Young, L.J. (2013). Intranasal oxytocin selectively attenuates rhesus monkeys’ attention to negative facial expressions. Psychoneuroendocrinology 38, 1748– 1756. 10.1016/j.psyneuen.2013.02.011.

74. Kanat, M., Heinrichs, M., Mader, I., Van Elst, L.T., and Domes, G. (2015). Oxytocin Modulates Amygdala Reactivity to Masked Fearful Eyes. Neuropsychopharmacol 40, 2632–2638. 10.1038/npp.2015.111.

75. Lieberz, J., Scheele, D., Spengler, F.B., Matheisen, T., Schneider, L., Stoffel-Wagner, B., Kinfe, T.M., and Hurlemann, R. (2020). Kinetics of oxytocin effects on amygdala and striatal reactivity vary between women and men. Neuropsychopharmacology 45, 1134–1140. 10.1038/s41386-019-0582-6.

76. Wahis, J., Baudon, A., Althammer, F., Kerspern, D., Goyon, S., Hagiwara, D., Lefevre, A., Barteczko, L., Boury-Jamot, B., Bellanger, B., et al. (2021). Astrocytes mediate the effect of oxytocin in the central amygdala on neuronal activity and affective states in rodents. Nat Neurosci 24, 529–541. 10.1038/s41593-021-00800-0.

77. Dumais, K.M., Alonso, A.G., Bredewold, R., and Veenema, A.H. (2016). Role of the oxytocin system in amygdala subregions in the regulation of social interest in male and female rats. Neuroscience 330, 138–149. 10.1016/j.neuroscience.2016.05.036.

78. Yao, S., Bergan, J., Lanjuin, A., and Dulac, C. (2017). Oxytocin signaling in the medial amygdala is required for sex discrimination of social cues. eLife 6, e31373. 10.7554/eLife.31373.

79. Djerdjaj, A., Rieger, N.S., Brady, B.H., Carey, B.N., Ng, A.J., and Christianson, J.P. (2023). Social affective behaviors among female rats involve the basolateral amygdala and insular cortex. PLoS ONE 18, e0281794. 10.1371/journal.pone.0281794.

80. Vörös, D., Kiss, O., Ollmann, T., Mintál, K., Péczely, L., Zagoracz, O., Kertes, E., Kállai, V., László, B.R., Berta, B., et al. (2023). Intraamygdaloid Oxytocin Increases Time Spent on Social Interaction in Valproate-Induced Autism Animal Model. Biomedicines 11, 1802. 10.3390/biomedicines11071802.

81. Bundzikova, J., Pirnik, Z., Zelena, D., Mikkelsen, J.D., and Kiss, A. (2008). Response of Substances Co-Expressed in Hypothalamic Magnocellular Neurons to Osmotic Challenges in Normal and Brattleboro Rats. Cell Mol Neurobiol 28, 1033–1047. 10.1007/s10571-008-9306-x.

82. Schulze, L., Renneberg, B., and Lobmaier, J.S. (2013). Gaze perception in social anxiety and social anxiety disorder. Front. Hum. Neurosci. 7. 10.3389/fnhum.2013.00872.

83. Tönsing, D., Schiller, B., Vehlen, A., Nickel, K., Van Elst, L.T., Domes, G., and Heinrichs, M. (2025). Altered interactive dynamics of gaze behavior during face-to-face interaction in autistic individuals: a dual eye-tracking study. Molecular Autism 16, 12. 10.1186/s13229-025-00645-5.

84. Lio, G., Corazzol, M., Fadda, R., Doneddu, G., and Sirigu, A. (2025). A neuronal marker of eye contact spontaneously activated in neurotypical subjects but not in autistic spectrum disorders. Cortex 183, 87–104. 10.1016/j.cortex.2024.10.022.

85. Gomez, J.L., Bonaventura, J., Lesniak, W., Mathews, W.B., Sysa-Shah, P., Rodriguez, L.A., Ellis, R.J., Richie, C.T., Harvey, B.K., Dannals, R.F., et al. (2017). Chemogenetics revealed: DREADD occupancy and activation via converted clozapine. Science 357, 503–507. 10.1126/science.aan2475.

